# Single-cell analysis of upper airway cells reveals host-viral dynamics in influenza infected adults

**DOI:** 10.1101/2020.04.15.042978

**Authors:** Yuming Cao, Zhiru Guo, Pranitha Vangala, Elisa Donnard, Ping Liu, Patrick McDonel, Jose Ordovas-Montanes, Alex K. Shalek, Robert W. Finberg, Jennifer P. Wang, Manuel Garber

## Abstract

Influenza virus infections are major causes of morbidity and mortality. Research using cultured cells, bulk tissue, and animal models cannot fully capture human disease dynamics. Many aspects of virus-host interactions in a natural setting remain unclear, including the specific cell types that are infected and how they and neighboring bystander cells contribute to the overall antiviral response. To address these questions, we performed single-cell RNA sequencing (scRNA-Seq) on cells from freshly collected nasal washes from healthy human donors and donors diagnosed with acute influenza during the 2017-18 season. We describe a previously uncharacterized goblet cell population, specific to infected individuals, with high expression of MHC class II genes. Furthermore, leveraging scRNA-Seq reads, we obtained deep viral genome coverage and developed a model to rigorously identify infected cells that detected influenza infection in all epithelial cell types and even some immune cells. Our data revealed that each donor was infected by a unique influenza variant and that each variant was separated by at least one unique non-synonymous difference. Our results demonstrate the power of massively-parallel scRNA-Seq to study viral variation, as well as host and viral transcriptional activity during human infection.

## Introduction

Influenza virus causes acute respiratory infections and results in 3-5 million cases of severe illness and up to 500,000 deaths worldwide annually (McMorrow et al., 2015; Paget et al., 2019; World Health Organization, 2014). Influenza virus primarily infects the human airway epithelium. Most infections are limited to the upper respiratory tract, which includes the nasal tract, sinuses, pharynx, and larynx, but infections can progress to the lower respiratory tract to cause severe disease (Iwasaki and Pillai, 2014). Investigations on host responses to influenza based on *in vitro* studies and *in vivo* murine models have provided insights in the viral pathology, but studies of host responses to influenza virus at the site of infection in humans are relatively sparse (Pulendran and Maddur, 2015). Human airway epithelium cell lines (e.g. A549) and air-liquid interface culture systems have been used to study the tissue responses to infection and to monitor viral life cycles (Slepushkin et al., 2001; Wu et al., 2016), but these systems cannot fully represent the complexity of infection. Similarly, animal models have been widely used in influenza virus research, yet interpretation of these results can be confounded by dissimilarities between these models and humans, and between lab-adapted and natural viruses (Hemmink et al., 2018; Hirst, 1947; Kim et al., 2015; Radigan et al., 2015; Shin et al., 2015). For example, laboratory mouse models such as C57BL/6J are not natural hosts to influenza virus, and human influenza virus inoculated in their nasal tract cannot progress to the lung (Ivinson et al., 2017). Similarly, the sialic acid glycosylation in murine airway cells is not representative of that found in humans (Ibricevic et al., 2006). Finally, host factors critical for controlling infection, such as MX1 which is nonfunctional in common laboratory mouse strains, differ between mice and humans (Shin et al., 2015; Verhelst et al., 2012). Indeed, studies of host-viral dynamics in the human airway *in vivo* are sparse (Pulendran and Maddur, 2015) and until recently, capturing the complexity of the human airway during infection has not been possible.

Single-cell RNA sequencing (scRNA-Seq) has been revolutionary in decoding processes in heterogeneous tissue. Studies on host tissue responses to influenza virus infection have been limited by bulk measurement that mask cell type-specific responses, changes that result from the asynchronous nature of infection, and changes in cell composition that occur during infection. scRNA-Seq permits simultaneously map human and viral transcripts in a single cell (Martin-Gayo et al., 2018; Russell et al., 2019b; Zanini et al., 2018a, 2018b), and has been applied to various host-virus interaction scenarios (Cristinelli and Ciuffi, 2018), including human immunodeficiency virus (HIV) (Bradley et al., 2018; Kazer et al., 2019) and hepatitis C virus (HCV) infection (McWilliam Leitch and McLauchlan, 2013). To date, scRNA-Seq in the context of influenza virus infection has been applied to the A549 cell line and mouse models (Kudo et al., 2019; Russell et al., 2018, 2019a; Steuerman et al., 2018; Vera et al., 2019). However, given that such studies cannot recapitulate the natural course of influenza infection, an investigation of the response to influenza virus infection and viral replication efficiency in its natural setting is warranted. To address these issues, we sought to obtain samples directly from humans with acute influenza infection for scRNA-Seq.

The nasal mucosa serves as a barrier for most airborne pathogens, and cells from this natural niche can reliably reflect host-pathogen interactions (Jochems et al., 2017). Therefore, we obtained nasal washes from influenza A virus (IAV) or influenza B virus (IBV)-infected adult donors presenting to our medical center to unbiasedly sample all cell types at the first line of defense against influenza virus infection. We sought to investigate the dynamics of influenza virus infection and the host response at the site of infection in humans at the molecular level, defining: 1) what cell types the virus infects in the human nasal tract, 2) the interhost variability of the virus strain, 3) the variability of inter-/intra-host virus expression patterns, and 4) the heterogeneity of host cell type responses to the viral infection. We used Seq-Well (Gierahn et al., 2017; Ordovas-Montanes et al., 2018), a portable microwell-based scRNA-Seq technology, to jointly define the infecting influenza virus and the host response at the molecular level. Our study defines diverse cell types that can be infected by and respond to the influenza virus infection in the human nasal tract, including a previously uncharacterized goblet cell population, specific to infected individuals, with high expression of MHC class II genes in infected individuals, and revealed the variability of viral sequences between infected individuals.

## Results

### Seventeen different cell types are captured in human nasal wash samples by scRNA-Seq

During the 2017-18 influenza season, we collected nasal washes from adults who presented to our medical center with influenza-like illness (ILI) and were diagnosed with either influenza A virus (IAV) or influenza B virus (IBV). We selected samples for further analysis after confirming the presence of viral transcripts by qPCR and moved forward with six IAV H3N2 and six IBV donor samples (see **Supplemental Table 1**). ILI activity was high at the time of collection, with prevalence of both IAV H3N2 and IBV Yamagata lineage (CDC, 2019). We also collected nasal washes from adult volunteers without ILI (n=6 donors). scRNA-Seq libraries were generated using the Seq-Well (Version 3) platform then sequenced (Gierahn et al., 2017) (**Figure 1A, Methods: Sample Collection, RNA sequencing**). In addition, for a subset of the donors, bulk RNA-Seq libraries were generated to confirm the presence of influenza virus. Additional details on the donors are available in **Supplemental Table 1**. Due to index swapping associated with the NovaSeq sequencing system (Costello et al., 2018) and a likely bead breakage of ChemGene beads on Drop-Seq based platforms, we carried out additional steps to process sequencing reads from scRNA-Seq and remove potential artifacts (**Methods: Special Notes, Figure S1A**).

**Figure 1.**
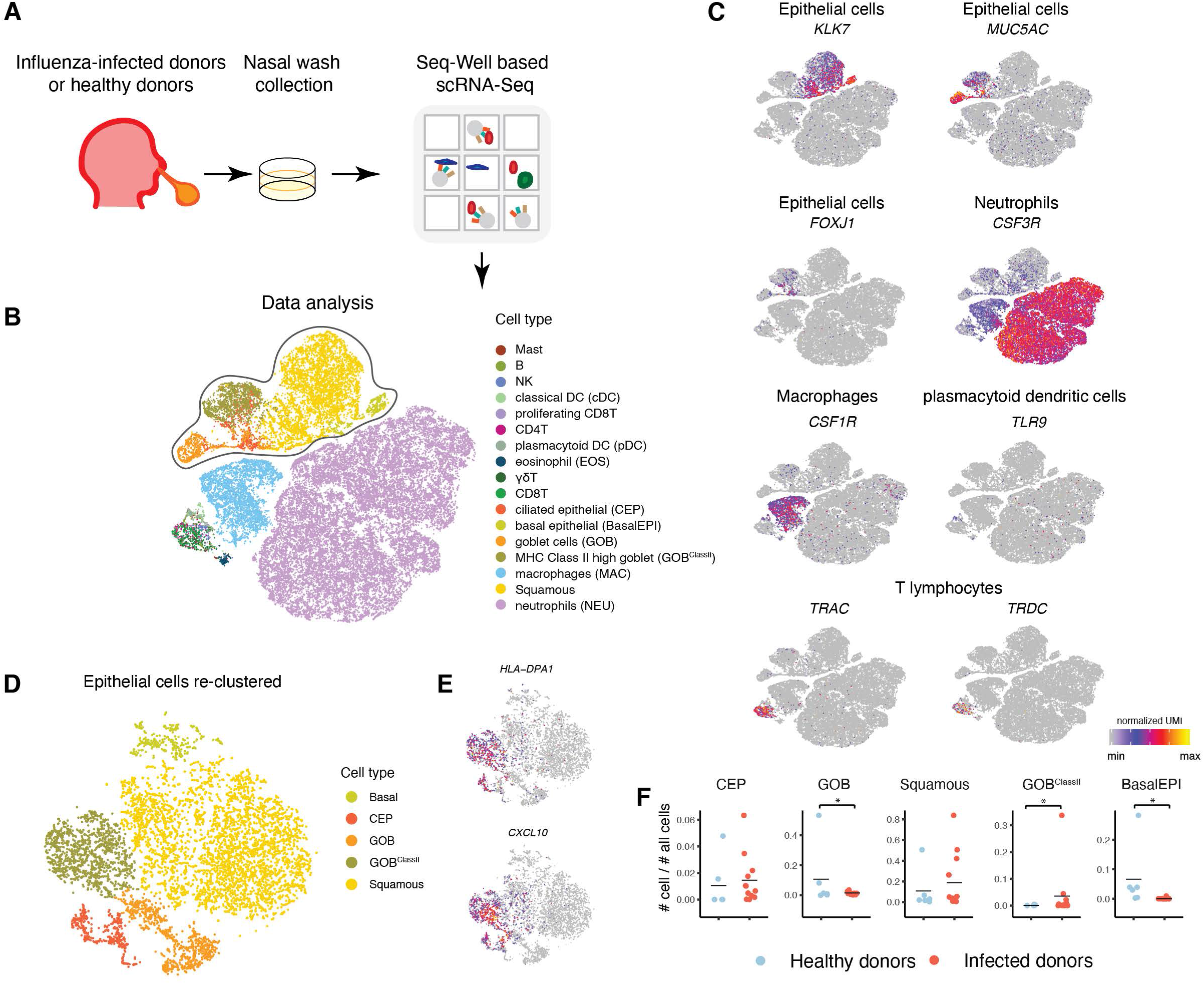
scRNA-Seq captures the cell type distribution in nasal wash samples from humans with acute influenza infection. **A.** Schematic of sample collection, processing, and data analysis. Donors had a diagnosis of influenza A virus (IAV) (n = 6) or influenza B virus (IBV) (n = 6) by rapid antigen test or respiratory virus panel and had detectable viruses in sequence or were healthy volunteers (n = 6). **B.** t-distributed stochastic neighbor embedding (tSNE) representation includes 35,480 cells clustering in two-dimensional space, colored by 17 distinct cell types identified. Cell barcodes with >1000 unique molecular identifiers (UMI) were denoted as cells. **C.** tSNE shows the normalized expression of gene markers for cell types found the major four broad clusters: an immune cell cluster including T lymphocytes (*TRAC, TRDC*) and dendritic cells (*TLR9* for plasmacytoid dendritic cells), epithelial cells (*FOXJ1, MUC5AC* and *KLK7*), macrophages (*CSF1R)*, and neutrophils (*CSF3R).* **D.** All epithelial cells were subsetted and reclustered. tSNE plot showing the new embedding of all epithelial cells with cells colored by cell types. **E**. The GOB_ClassII_ cell cluster shows high expression of HLA class II transcripts and interferon response genes (*HLA-DPA1* and *CXCL10* shown as examples). **F.** Fraction of each epithelial cell type out of all cells found in each donor. The number of each cell type from each healthy (n=6) and infected (n=12) donor was plotted here. The black line shows the mean. *: p < 0.05, Wald test.

We obtained a total of 35,480 cells across all samples (n = 18), in which we detected 18,870 genes. We carried out a two-step clustering approach to identify cell types present in the scRNA-Seq data. In the first round we identified four broad clusters (**Figure 1B**), including neutrophils (shown in light purple), which have been largely absent in data obtained from other scRNA-Seq technologies (Smillie et al., 2019). The remaining cells include macrophages (light blue), epithelial cells (circled), and leukocytes such as lymphocytes and dendritic cells (lower left). Conventional markers for select cell types, namely *KLK7*, *MUC5AC*, and *FOXJ1* for epithelial cells (McCauley et al., 2018), *CSF3R* for neutrophils (Ancuta et al., 2009), *CSF1R* for macrophages (Lavin et al., 2015), *TLR9* for plasmacytoid dendritic cells, and *TRAC and TRDC* for T lymphocytes, further delineated cell heterogeneity inside each broad cluster (**Figure 1B-C**).

Next we independently analyzed each of the four broad clusters (**Methods: Clustering**) in order to distinguish additional cell populations. Subclustering of these initial broad clusters revealed all of the known cell types in the upper respiratory tract (Denney and Ho, 2018; Deprez et al., 2019; Ruiz García et al., 2019; Vareille et al., 2011; Wu et al., 2016). The clusters dominated by neutrophils (NEU, *CSF3R*+) and by macrophages (MAC, *CSF1R*+) showed homogeneous populations that did not cluster into subtypes, however in both clusters cells from infected donors showed an increased activation state as suggested by high expression of *ISG15, RSAD2* and *IFIT3* (**Figure 1B-C, Figure S1B-D**). Epithelial cells were subclassified into non-cycling basal epithelial cells (BasalEPI, *KRT5*+), squamous cells (*KLK7*+) (Pettus et al., 2009), goblet cells (GOB, *MUC5AC+),* ciliated epithelial cells (CEP, *FOXJ1*+) (Ruiz García et al., 2019), and a cluster of class II HLA+ highly expressing cells that also expressed Goblet marker genes *(MUC5AC* and *MUC16)* (GOB_ClassII_) (**Figure 1D, Figure S2C**). GOB_ClassII_ cells also expressed specific interferon-stimulated genes at high levels (e.g. *CXCL10, CXCL11, IFI6,* and *IFITM3*) **(Figure 1E, Figure S2C)**. Within the ciliated epithelial cell cluster, we found a subset of cells with high viral mRNA expression (**Figure S2D**). Closer inspection showed these cells are a mixture of *FOXJ1*+ cells (n = 115), *MUC5AC*+ cells (n = 16), *FOXJ1+ MUC5AC*+ double positive cells (n = 23), and *FOXJ1-MUC5AC-* double negative cells (n = 156) (**Figure S2D,F**). We refer to these cells generically as CEP.

Leukocyte subclustering revealed the presence of CD8+ T cells (CD8T, *CD8A*+), proliferating CD8+ T cells (*MKI67*+), gamma delta T cells (γδT, *TRDC+, TRGC1/2*+), CD4+ T cells (CD4T, *TRAC+, CTLA4*+), natural killer cells (NK, *FCGR3A+, NKG7*+), and B cells (B, *CD19*+); this same cluster included plasmacytoid dendritic cells (pDCs, *TLR7/9/10*+) and conventional dendritic cells (cDCs, *CLEC10A*+) (Li et al., 2020). We also found two types of granulocytes in this cluster: eosinophils (EOS, *CLC*+) (Gomolin et al., 1993) and mast cells (MAST, *HDC*+) (Li et al., 2018) (**Figure S3)**.

In total, we identified 17 distinct cell types, including five epithelial cell types (CEP, GOB, BasalEPI, Squamous and GOB_ClassII_), six myeloid cell types (NEU, EOS, MAST, MAC, cDC and pDC), and six lymphoid cell types (CD4T, CD8T, proliferating CD8T, γδT, B, and NK) **(Figure 1B, Figure S1E-F).** Marker genes for each cell type are provided in **Supplemental Table 4.**

### A goblet cell type expressing high levels of MHC class II is abundant in the infected upper respiratory tract

Acute and chronic respiratory diseases harm mucosal barrier integrity, affecting cell type composition by inducing epithelium regeneration and immune cell infiltration (Ordovas-Montanes et al., 2018; Ruiz García et al., 2019; Vieira Braga et al., 2019). We investigated whether the distribution of cell types sampled in infected donors differs from that of healthy controls. Overall, neutrophils and all five types of epithelial cells are more abundant in infected donor samples (21,727 neutrophils, 8,938 epithelial cells), after taking into account the randomness of sampling and the high variability in the number of cells found in each sample (Xu et al., 2019) (**Methods: Cell type enrichment statistical test**). Infected donors generally yield more cells than healthy donors (t-test, *P* = 0.003, **Figure S1G**). Although the fraction of epithelial cells was variable across donors (**Figure S1E-F**), infected donors yielded more GOB_ClassII_ compared to healthy donors (p = 0.03, Wald test), with 3 infected donors having more than 2% of their cells classified as GOB_ClassII_. On the other hand, healthy donors contributed more BasalEPIs (p = 0.015, Wald test) and GOBs (p = 0.015, Wald test) (**Figure 1F**). Infected donors also yielded more innate immune cells and adaptive immune cells, such as MAC, cDC, proliferating CD8T cells and *γδ*T cells (p = 0.015, p = 0.03, p = 0.0003, and p = 0.03, respectively, Wald test) (**Figure S1H**). Eosinophils were found in only seven infected donors (six of them are IAV donors with 91% of EOS derived from donor IAV7).

Taken together, the combined scRNA-Seq data generated from human nasal wash samples contains cell types that are both normally resident in the upper respiratory tract as well as induced or recruited during the viral insult. The various types of epithelial cells found in these data, and the GOB_ClassII_, in particular, which we specifically identified in infected donors, reflect the important cellular dynamics at play during influenza virus infection.

### Viral transcripts are detected in epithelial cells as well as some immune cells

Influenza virus predominantly infects epithelial cells but can also infect various types of immune cells (Iwasaki and Pillai, 2014). In a mouse model, viral mRNA was detected in all lung cell types sampled, with epithelial cells harboring the highest viral loads (Steuerman et al., 2018). Furthermore, *in vitro* and *in vivo* experiments have shown a wide range of influenza viral transcriptional activity across infected cells (Russell et al., 2018; Steuerman et al., 2018). Here, we sought to both identify virally infected cell types and determine the cellular viral load in human nasal wash samples.

Our *in vivo* samples revealed a wide range of viral expression even within a given cell type (**Figure 2A**). However, mere detection of low levels of viral mRNA does not provide certainty of viral infection since these mRNAs may originate from ambient RNAs during scRNA-Seq sample preparation (Lun et al., 2019; Steuerman et al., 2018; Young and Behjati, 2018). Ambient RNAs from lysed cells can result in a sample-specific contamination. In particular, virus-induced cell lysis causes viral RNAs to dominate the ambient RNA pool. We therefore estimated a sample-specific distribution of ambient influenza mRNA contamination and predicted cells most likely to be infected using a hurdle zero inflated negative binomial (ZINB) model and a support vector machine (SVM) classifier (**Figure 2B, Figure S4, Methods: Identifying viral infected cells)**. Briefly, cells were first grouped into healthy, virus-, and virus+ based on viral exposure. A ZINB model was applied for virus+ and virus- cells in each highly infected donor, and predicted if a cell was infected or not. Based on the genes that are significantly differentially expressed between predicted infected cells and predicted uninfected virus- cells, a SVM classifier was built for selected cell types (CEP and Squamous cells) to classify infected or uninfected (bystander) cells (workflow illustrated in **Figure 2B**). A total of 671 out of 30,546 cells (2.2%) from the infected donors were classified as infected (**Figure 2C**). The remaining cells from infected donors were classified as bystander cells, and cells from healthy donors were termed healthy cells. Viral mRNA expression was robustly detected in 11 out of the 17 cell types identified in infected donors (**Figure S4E**). We found that all subtypes of epithelial cells were infected, as well as NEU, MAC and T lymphocytes. However, no dendritic cells, NK, B, or mast cells were predicted to be infected according to the ZINB model.

**Figure 2.**
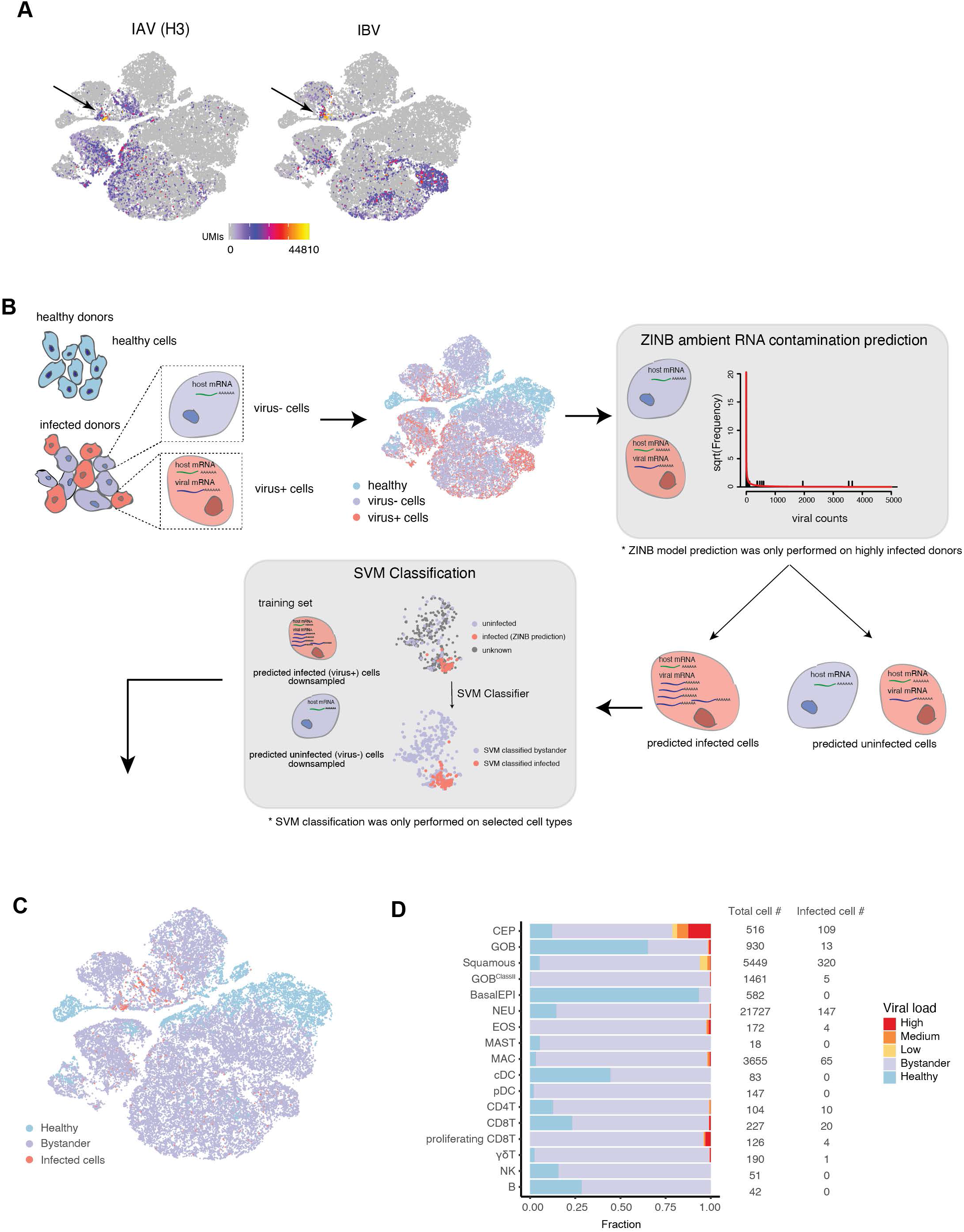
Viral transcripts are detected in epithelial cells as well as immune cells in human nasal washes. **A.** tSNE representation shows the raw expression of IAV (H3N2) or IBV transcripts in all cells before identifying infected cells. Viral metagenes, composed of all eight segments, are plotted here. The arrows point to the group of highly infected cells. **B.** Workflow of identifying viral infected cells from infected donors. This method is described in detail in **Methods: Identifying Viral Infected Cells. C.** tSNE representation of cells in different cellular states: healthy, bystander, and infected after classification. **D.** The distribution of viral load in cell types predicted to be infected by the ZINB model and the SVM classifier. Viral load for each cell is calculated based on the fraction of viral transcripts to total transcripts. Viral load status is empirically classified by bystander (cells predicted and classified as bystander by ZINB model and SVM classifier), low, medium, or high based on the tertile of viral gene expression rank in a cell.

Based on the expression rank of viral genes, we divided the expression rank into tertiles and classified the viral infection states of each infected cell as low, medium or high (**Figure 2D**). This stratification revealed that ciliated epithelial cells carry the largest viral load (**Figure 2D**) for cells from both IAV- and IBV-infected donors. (**Figure S5**).

### M and NP segments are the most highly expressed, while the NS segment drops out in highly infected cells

IAV and IBV genomes each contain eight segments, with some segments encoding more than one protein through alternative splicing, some of which disrupt interferon production of the host cell (Garfinkel and Katze, 1993; Kochs et al., 2007a). Success in expressing all eight influenza genome segments in a cell is critical for reproductive infection for both IAV and IBV (Sheng et al., 2018; Vasilijevic et al., 2017). An expression hierarchy of M>NS>>NP>NA>HA>>PB2∼PB1∼PA was reported by previous *in vitro* single cell transcriptome studies in IAV-infected A549 cells and by bulk measurements of IAV-infected MDCK cells (Hatada et al., 1989; Russell et al., 2018). It is unclear whether this expression pattern holds true *in vivo*. Viruses lacking one or more segments have a higher chance of being recognized by the host (Vasilijevic et al., 2017). IAV deficient in the segment NS has been associated with higher antiviral response *in vitro* (Russell et al., 2019a). The NS segment encodes for the NS1 protein, which antagonizes the host antiviral response by interfering with IRF3 to suppress interferon (IFN) and tumor necrosis factor (TNF) expression (Geiss et al., 2002; Kochs et al., 2007b).

To address whether such phenomena occur *in vivo* during human infection, we examined viral transcripts in samples from influenza-infected donors. First, we examined expression hierarchy. Because certain influenza genome segments yield transcripts with the same 3’-end sequence by alternative splicing (Dubois et al., 2014), and our data captures viral mRNA transcripts with a bias toward the 3’-end, we were unable to differentiate those alternatively spliced transcripts. While targeted probes could be used to specifically measure expression levels of different isoforms in future studies, here, we quantified expression at the segment level. In cells with high viral loads, we detected expression of all eight segments in the majority of infected cells (IAV: 61%, IBV: 90%) (**Figure 3A**). Contrary to *in vitro* studies, we observed that segment 5 (NP) and segment 7 (M) were most highly expressed in both IAV- and IBV-infected cells, regardless of cell type, while the other six segments had lower expression (**Figure 3B**). The trend was consistent across cells with lower infection levels, and for both IAV and IBV samples (**Figure 3B**).

**Figure 3.**
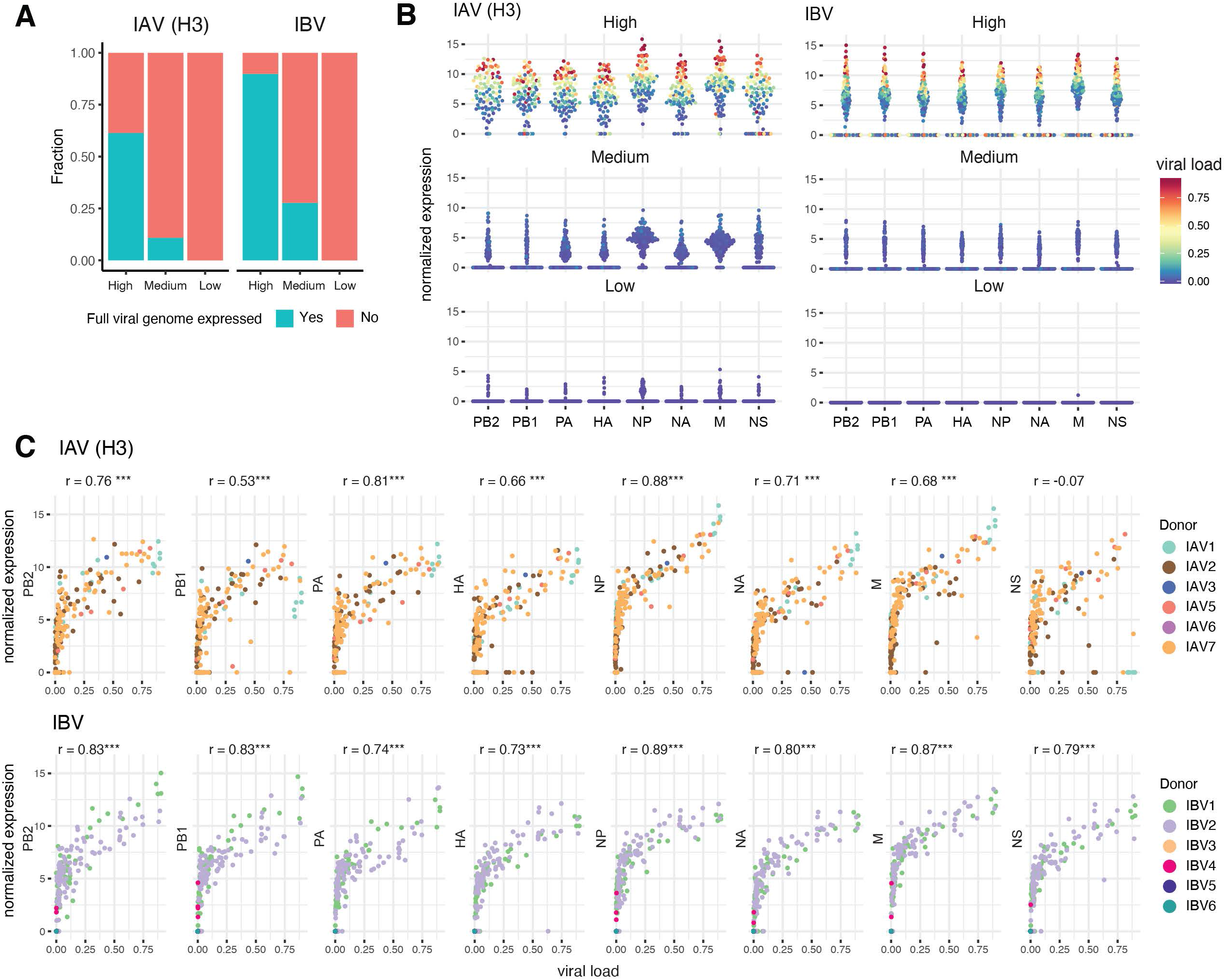
The M and NP segments are the most highly expressed, while the NS segment drops out in highly infected cells. **A.** The fraction of cells either expressing all eight viral segments or having at least one segmental dropout in each viral load state. **B.** Normalized expression levels of each viral segment from either IAV- or IBV-infected cells. Viral segment expression is colored by the viral load of their host cell. **C.** Scatter plots showing the correlation between viral load (the fraction of viral mRNA from the virus for each cell) and the normalized expression level for each viral segment in each infected cell per donor. The Pearson correlation coefficient r is calculated for each segment. ***: P < 0.0001.

Next, we considered viral segmental dropout. For all IBV segments and for all but one IAV segment (segment 8, NS), a clear correlation was observed between viral load and segment expression (r = 0.53 - 0.89, Pearson’s correlation). However, the IAV NS segment had a poor correlation with viral load (r = −0.07, Pearson’s correlation), especially in cells with the highest viral load. In fact, for 7 out of the 11 cells with the highest viral loads (viral load > 75%), NS was not detected (**Figure 3C**). This drop-out would not be otherwise expected assuming that segment detection is similar for all segments (P < 0.0001, Chi-squared test) (**Figure 3C**). IAV segment expression in each infected cell also had higher variability compared to IBV segment expression (**Figure S6**). However, limited by the small number of NS-negative cells, we were unable to conclude on the difference between IFN production efficiency between NS-positive and NS-negative cells.

### scRNA-Seq allows detection of single nucleotide variants (SNVs) in viral transcripts and shows that at least one unique strain infected each individual

Influenza viruses possess high evolutionary potential due to their high mutation rate (Chen and Holmes, 2006; Hadfield et al., 2018; Nobusawa and Sato, 2006). However, it is unclear how different selection pressures and stochastic processes affect the overall viral evolution dynamics (McCrone et al., 2018). A number of studies have investigated the inter- and intra-host diversity of the viral sequences in virus-challenged humans and in animal models (Iqbal et al., 2009; Leonard et al., 2016; Lin et al., 2019; Murcia et al., 2010, 2012). These studies, performed in a laboratory-controlled setting, highlight the rapidness of allele fixation within the host and the genetic diversity between hosts of influenza viruses. Because the evolutionary dynamics between hosts and within hosts during the course of natural infections in humans is less clear, we looked for single nucleotide variants (SNVs) in our dataset.

By combining all scRNA-Seq influenza mapping reads for each donor we obtained deep coverage of the influenza genome (average 595x per donor, **Figure S7**). Since scRNA reads are associated with technical biases (McCrone and Lauring, 2016) and index swapping that occurred during the sequencing of pooled variables introduced additional variability (Costello et al., 2018), we restricted our analysis to the most common SNVs and implemented many steps to remove known biases that can affect variability estimates (**Methods: Virus genotyping and SNV calling**). We then estimated influenza sequence variability from the samples collected in a single-season of geographically localized infection, by using available sequences for IAV (H3N2) (NCBI:txid2069524) and IBV (NCBI:txid2067645) strains collected in Massachusetts earlier in the same season **(Method: Genome Alignment**). For each infected donor, we used stringent criteria to identify positions that differed from the reference. We excluded two IBV donors (IBV3 and IBV5) due to low infection levels that resulted in low read coverage. We built consensus sequences for IAV and IBV, and identified positions that harbored SNVs in our patient samples. We found a total of 80 SNVs in the six donors with IAV, and 39 SNVs in the four donors with IBV (**Supplemental Table 2**). Bulk RNA-Seq libraries generated from excess nasal cells not loaded on Seq-Well confirmed the variant calls (with the exception of IAV6 and IAV7, for which no extra cells were available for bulk RNA sequencing). These high confidence SNVs were used to estimate interhost variability. For each donor, we estimated the major SNV for each position as well as its frequency **(Methods: Virus genotyping and SNV calling**). We find that each donor has at least one unique SNV, and IAV and IBV had an average of 6.5 unique and 7.5 unique SNVs, respectively (**Figure 4A-B**). Parsimony analysis of the variable positions showed no correlation with the timing of the sample collected (**Figure 4C, Supplemental Table 1**).

**Figure 4.**
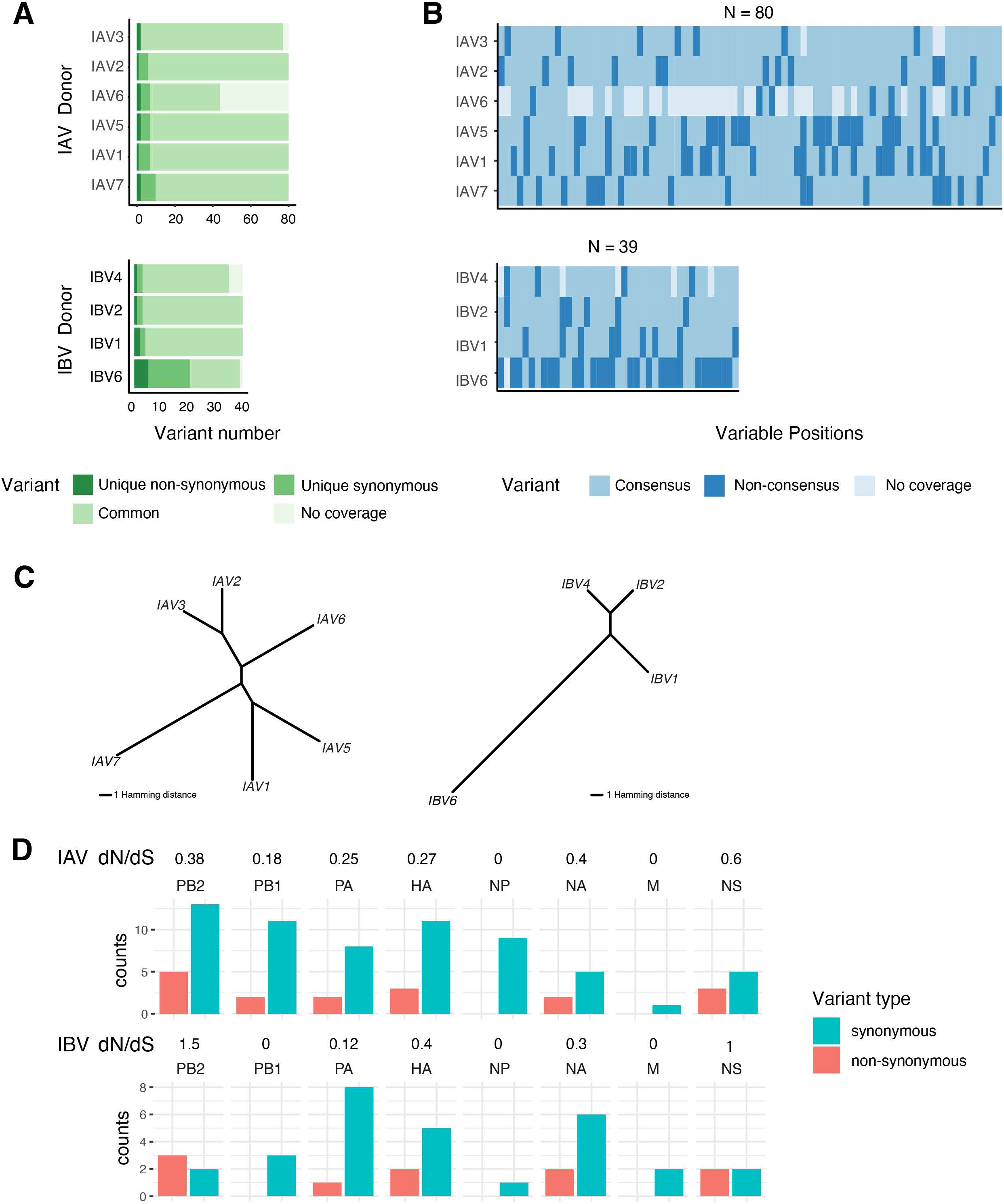
scRNA-Seq allows detection of SNVs in viral transcripts and shows that at least one unique strain infected each individual. Sequencing reads mapped to influenza reference genomes from valid cells (>1000 UMIs) were used to identify SNVs present in each donor. Variant calls were made based on the reference genomes, and were converted to consensus if present in more than half of donors for each viral subtype to avoid reference bias. **A.** Quantification of SNVs private to each donor. IBV3 and IBV5 are not shown in this analysis due to lack of enough infected cells. Of all variants found in each viral subtype (n=80 for IAV, n=39 for IBV), each donor harbored unique SNVs. Variable positions not covered in valid cells are marked as no coverage. These positions are not considered in parsimony analysis. **B.** Barcode plot showing the classification of each variable position found in each donor. **C.** Unrooted dendrogram showing the SNVs Hamming distances among donors. The donor numbering represents the sequence of collection time. No correlation between sample collection time and position on the dendrogram is found. **D.** The number of non-synonymous and synonymous variants from all SNVs defined (n=80 for IAV, n=39 for IBV). Variant positions with no coverage in any of the donors are not considered. The dN/dS ratio (ratio of the number of non-synonymous SNVs to synonymous SNVs) is noted on top of each segment.

We found non-synonymous and synonymous SNVs in both IAV and IBV. IAV and IBV genomes showed a similar non-synonymous to synonymous ratio (dN/dS) with IAV having a dN/dS = 0.25 and IBV dN/dS = 0.34 (**Figure 4D**). Regardless of viral strain, non-synonymous SNVs in segments 5 and 7 (NP and M) are consistently absent. We observed at least one unique non-synonymous SNV in each donor’s viral genomes (**Figure 4A**). For IAV, non-synonymous SNVs that may affect protein charge, hydrophobicity or structure are as follows: PB2: T64I and A707S, PB1-F2: G23E; HA: K43E and T228A; NA: S217L and Y284H; NS: M119V, and F128S. For IBV, those SNVs are PB2: S591R; PA: T60A; NA: L74P and G234R; NS: N199D and E235K (all non-synonymous SNVs are provided in **Supplemental Table 3**). Altogether, each IAV or IBV infected donor has a distinct viral strain, harboring unique SNVs that are not present in other donors, and each donor had at least one non-synonymous SNVs. This result indicates that even in a relatively constrained geographic location, thousands of different influenza viral strains are likely circulating during a season.

### Viral infection induces type I and type III IFN signaling in infected cells and IFN signaling responses from bystander cells

We next examined differential gene expression between all bystander and infected epithelial cells from donors with IAV or IBV and healthy epithelial cells from control donors. Irrespective of the viral type, 225 genes were found to be upregulated in infected epithelial cells compared to bystander cells, and 376 genes were found to be uniquely upregulated in bystander cells compared to healthy controls (**Supplemental Table 5**). Differential expression analysis revealed the upregulation of type III IFNs (*IFNL1-3*) in infected epithelial cells (**Figure 5A**). As type III IFNs are critical in barrier tissue response to infection (Broggi et al., 2020), epithelial cells could be the major producers of type III IFN in the human nasal tract during influenza infection. Inspection of type I and type III IFN transcripts across all cell types showed that indeed, infected epithelial cells are the main producers of both type I IFN and type III IFN transcripts (**Figure 5B-C**). Infected CEPs dominated the *IFNB1* and type III IFN transcript production (**Figure 5D**). Type I and III IFN transcript upregulation was also seen in select immune cells during infection (**Figure 5B-D**).

**Figure 5.**
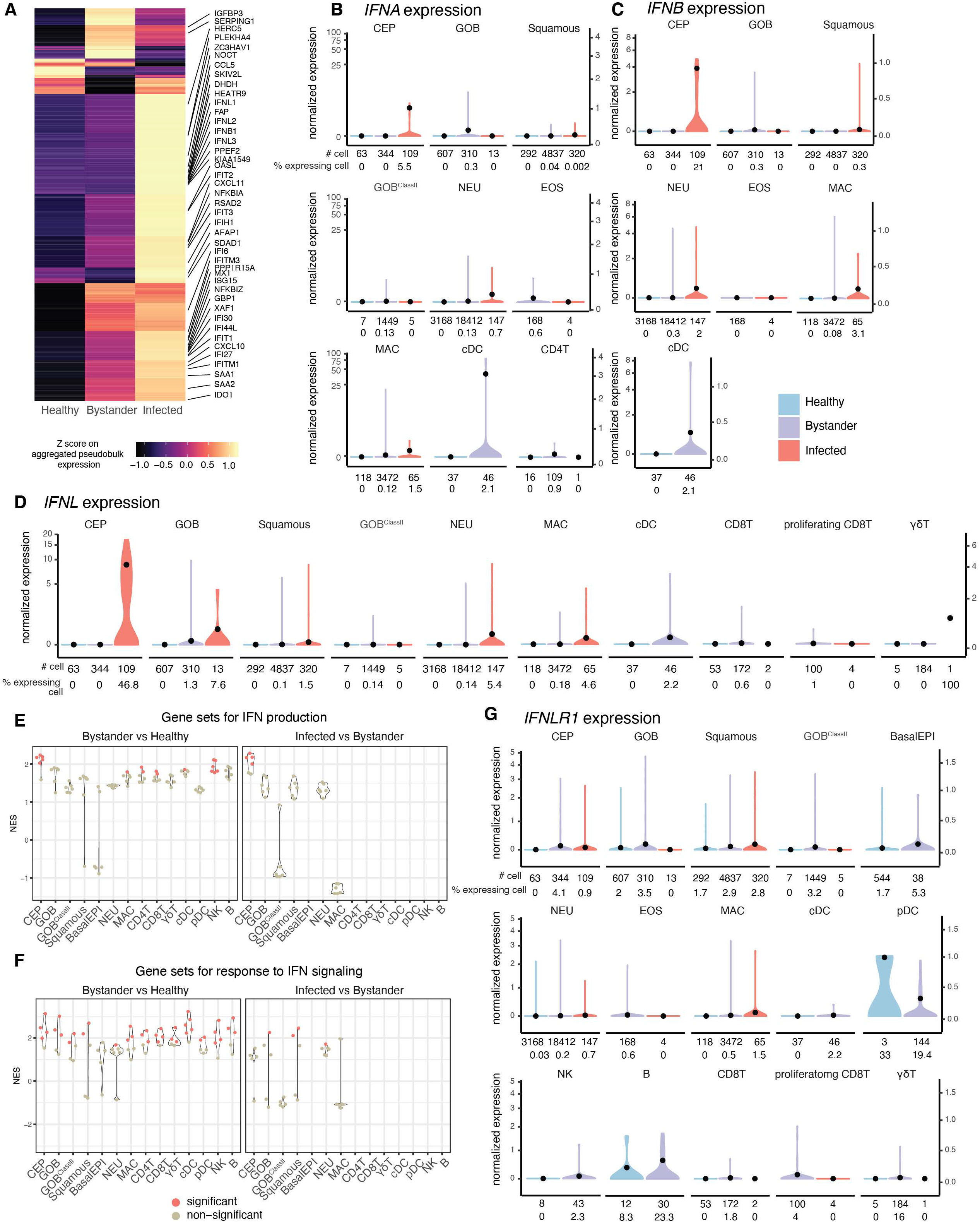
Viral infection induces type I and type III IFN production in infected cells and IFN signaling response from bystander cells. *IFNA* and *IFNL* metagene expressions are calculated as the sum of all mapped *IFNA* genes and mapped *IFNL* genes. Only cell types that have non-zero expression of each metagene are shown here. Gene expression counts shown in this figure are normalized counts. **A.** Heatmap on the row-scaled normalized expression z-scores of all significantly differentially expressed genes (DEGs) identified in epithelial bystander cells in comparison to healthy cells and infected cells in comparison to bystander cells. Aggregate pseudobulk gene expression (see **Method: Aggregated pseudobulk gene expression calculation**) was calculated per cellular infection state of all epithelial cells. The top 40 DEGs are annotated with text on the heatmap. **B.***IFNA* metagene expression levels in different cellular states across cell types. The black points are the mean expression levels. **C.***IFNB1* expression levels in different cellular states across cell types. The black points are the mean expression levels. **D**. *IFNL* metagene expression levels in different cellular states across cell types. The black points are the mean expression levels. **E.** The enrichment of type I IFN production gene sets (GO:0032648, GO:0032481, GO:0032728, GO:0032727, GO:0032647, GO:0032479) tested by GSEA. Enrichment of gene sets was tested between bystander cells and healthy cells, and between infected cells and bystander cells for each cell type. The normalized enrichment score (NES) for each comparison per cell type is shown here. Positive NES denotes enrichment of gene sets in bystander cells when compared to healthy cells, and in infected cells when compared to bystander cells. Gene sets were not tested in cell types with less than 10 cells in any cell state, thus left blank on the plot. NES values are colored by their significance. **F.** The enrichment of type I IFN response gene sets (GO:0034340, GO:0035456, GO:0035455, GO:0060338, GO:0035458, GO:0005132) tested by GSEA. **G**. *IFNLR1* expression levels in different cellular states across cell types. The black points are the mean expression levels.

Given the clear upregulation of classical IFN response genes (e.g. *CCL5, IFIH1, IFIT1-3, IFI6,27,30*) in both bystander and infected cells, with a stronger upregulation in infected cells (**Figure 5A**), we sought to better understand how cells respond to IFNs. To this end, we compiled “IFN production” and “IFN response” gene sets, and performed gene set enrichment analysis (GSEA) (Subramanian et al., 2005) (**Figure 5E-F, Supplemental Table 6**). Consistent with the expression of the *IFNL* cytokine, the IFN production gene sets were most highly upregulated in bystander and in infected CEPs. IFN response gene (IRG) sets, on the other hand, revealed an interesting pattern: while IRGs were induced in nearly all bystander cells relative to healthy cells, especially CEPs, NEUs did not exhibit this trend (**Figure 5F, left panel**). This was despite the fact that NEU were the most abundant immune cell in our samples and hence the cell type for which we had most power to detect such trend. Beside bystander NEUs, we also found that infected CEPs also failed to upregulate IRGs. This was surprising given that infected CEPs had the highest levels of *IFNLs* (**Figure 5F, right panel**). IFN unresponsiveness within infected CEPs may reflect the ability of influenza virus to block IRGs, but not the production of *IFNLs*. Furthermore, inspection of transcript abundance for type I receptors (*IFNAR1* and *IFNAR2*) and type III IFN receptors (*IFNLR1* and *IL10RB*), revealed that while *IFNAR1, IFNAR2* and *IL10RB* transcripts were ubiquitous (**Figure S8A-C**), expression of *IFNLR1* (encoding a member of the type III IFN receptor heterodimer), was cell type-specific with the lowest expression found in NEU, the cell type with the lowest IRG response (**Figure 5G**). This suggested that the response to influenza detection was driven by type III IFNs, which were produced in infected cells, specifically infected CEPs. Type III IFNs likely triggered a cell type-specific response in cells having robust expression of *IFNLR1*. The low expression of *IFNLR1* in neutrophils also indicated that the responsiveness of neutrophils to type III IFN signaling was context-dependent, as a high expression of *IFNLR1* was reported in neutrophils obtained from bone marrow and blood (Broggi et al., 2017; Espinosa et al., 2017; Galani et al., 2017).

Given that MHC class II upregulation by IFN-γ has been shown in A549 cells (Uetani et al., 2008), the induction of MHC class II transcripts observed in goblet cells (GOB_ClassII_) during influenza infection was likely a consequence of IFN response activity. In fact, we found that MHC class II gene expression was higher in bystander and infected cells compared to healthy, influenza-naive cells for most epithelial cell types (**Figure S9**), suggesting that epithelial cells in general have the potential to become antigen presenting cells under pathogen attack, with goblet cells exhibiting the highest potential.

## Discussion

In this study, we generated the first comprehensive cell atlas of human nasal cells during influenza infection. Our study uniquely captured the following: (1) cell populations both harboring virus and responding to infection in the natural infection niche, (2) computational methods for identifying virally infected cells and viral SNVs from scRNA-Seq data, (3) diversity of viral sequences for seasonal influenza viruses within a proximal geographical distance, (4) human transcriptomic modulation in individual cells from the site of primary influenza viral infection, and (5) type III IFN as the predominant response to influenza infection in human nasal tract.

Our data revealed two strong trends regarding epithelial cells. On one hand, MHC class II gene expression in epithelial cells was overall upregulated. Most strikingly, a subset of goblet-like cells expressed high levels of transcripts from MHC class II genes. This suggested that the epithelium may complement the professional antigen presentation cells at the site of infection. Based on *in vitro* studies and studies on polyps from patients with allergic diseases (Arebro et al., 2016; Kalb et al., 1991; Salik et al., 1999; Wang et al., 1997), increased levels of MHC class II transcripts in epithelial cells may be a standard response to IFNs. On the other hand, basal epithelial cells were depleted in the upper respiratory tract. Tissue damage caused by the influenza virus infection may drive basal epithelial cells to differentiate for tissue repair, making basal cells less likely to be detected by transcriptional analyses. However, we cannot rule out the possibility that our observation is affected by the sampling bias, where infected donors shed more fully differentiated epithelial cells.

We identified a large number of neutrophils in our nasal wash populations, and they were clearly split into two cell populations, one of which expressed active markers (e.g. *ISG15, IFIT3,* and *RSAD2*) mostly from infected donors. Neutrophils are mostly depleted in scRNA-Seq datasets generated from the 10x Genomics platform (Park et al., 2018; Smillie et al., 2019). Neutrophils are prone to activate their RNases that prevent efficient mRNA capture during prolonged incubation time and harsh conditions during single-cell capturing in technologies such as 10x and inDrop (Schwartz et al., 2018). We found eosinophils, another important granulocyte, predominantly in one patient. It is unclear what role, if any, eosinophils have in combating influenza infection. The individual where we found eosinophils may have been afflicted with an allergic reaction in addition to an influenza infection. Further studies with larger sample sizes will provide insight to this matter.

Studies in human primary polarized airway epithelial cell cultures have suggested that airway epithelial cells are primarily responsible for type III IFN production (Fox et al., 2015; Ioannidis et al., 2013; Killip et al., 2015; Klinkhammer et al., 2018; Okabayashi et al., 2011). We found that infected ciliated epithelial cells from human nasal cells were the major producers of type III IFN transcripts during natural infection, within days following symptoms and diagnosis, consistent with the observations from *in vivo* models and air-liquid interface systems. Interestingly, we noticed that viral infection seems to disrupt the IFN response in infected ciliated epithelial cells, as evidenced by downregulation of interferon response genes compared to bystander cells. However, epithelial cells maintained high expression of genes in the IFN production pathway. We could not however evaluate whether IFNs themselves were suppressed perhaps due to the small size of IFN genes that reduces their detection rate in scRNA-Seq libraries. Taken together the effects on IRGs show a failure to suppress viral infection by highly infected epithelial cells.

We developed two important methodologies that can be applied to other host-virus dual transcriptome scRNA-Seq studies: 1) a statistical test for identifying viral infected cells from scRNA-Seq data, overcoming the confounding ambient RNA contamination in scRNA-Seq, and 2) an approach that uses scRNA-Seq to map virus sequence variability in addition to viral gene expression. In fact, despite limitations from the 3’-end transcript bias of scRNA-Seq, we showed that we were able to define viral sequences and make viral SNV calls for each highly infected donor. Although we only captured viral transcripts (instead of the viral genome), our stringent SNV calling method allowed us to identify viral genomic variations and differentiate them from random errors introduced by RNA polymerases. We showed that, within a small geographical region and a defined sampling time frame, influenza viruses from infected donors have variabilities in their genetic sequences, with each donor harboring a unique viral sequence. Twenty percent of these variants also had predicted amino acid changes. The fact that we are able to identify SNVs from scRNA-Seq data opens up the possibility to directly study the viral sequence in each single cell from primary human samples, and suggests that these techniques can be similarly applied to other viruses. Because we were limited by the short read lengths and the index swapping issues that particularly confound low frequency variant calls (**Methods**: **Virus genotyping and SNV calling**) (Costello et al., 2018), we were unable to characterize the intrahost viral genetic variability. However, these two issues can be addressed in the future by using a different sequencing strategy that reduces index swapping, and using longer reads that capture a greater fraction of each transcript, doing so should empower follow up studies to explore intrahost variability of the viral sequence.

Our study clearly showed the power of studying upper respiratory tract samples from influenza-infected humans coupled with single-cell technologies to understand local host responses. Future studies applied to other human respiratory viruses will allow for comparative analysis of the host-pathogen interplay and potential identification of host or viral factors that are responsible for increased or reduced virulence (Ziegler et al., 2020). Technological advances that bring sequencing applications closer to patient care will delineate the mechanisms that mediate influenza pathogenesis in humans, which will be crucial for designing improved vaccines and therapeutics against influenza and other respiratory viruses.

## Supplemental Tables and Figure Legends

***Supplemental Table 1***

**Summary of donor information, scRNA-Seq and bulk RNA-Seq library preparation and viral detection.**

***Supplemental Table 2***

**Details on viral genetic variation detection.**

***Supplemental Table 3***

**Details on non-synonymous variants of the viral genomes.**

***Supplemental Table 4***

**Top 50 differentially expressed gene markers for each cell type tested by *edgeR***.

***Supplemental Table 5***

**edgeR results of bystander cells compared to healthy cells and infected cells compared to bystander cells in epithelial cells.**

***Supplemental Table 6***

**Genes from either IFN production gene sets or IFN response gene sets**

**Figure S1.**
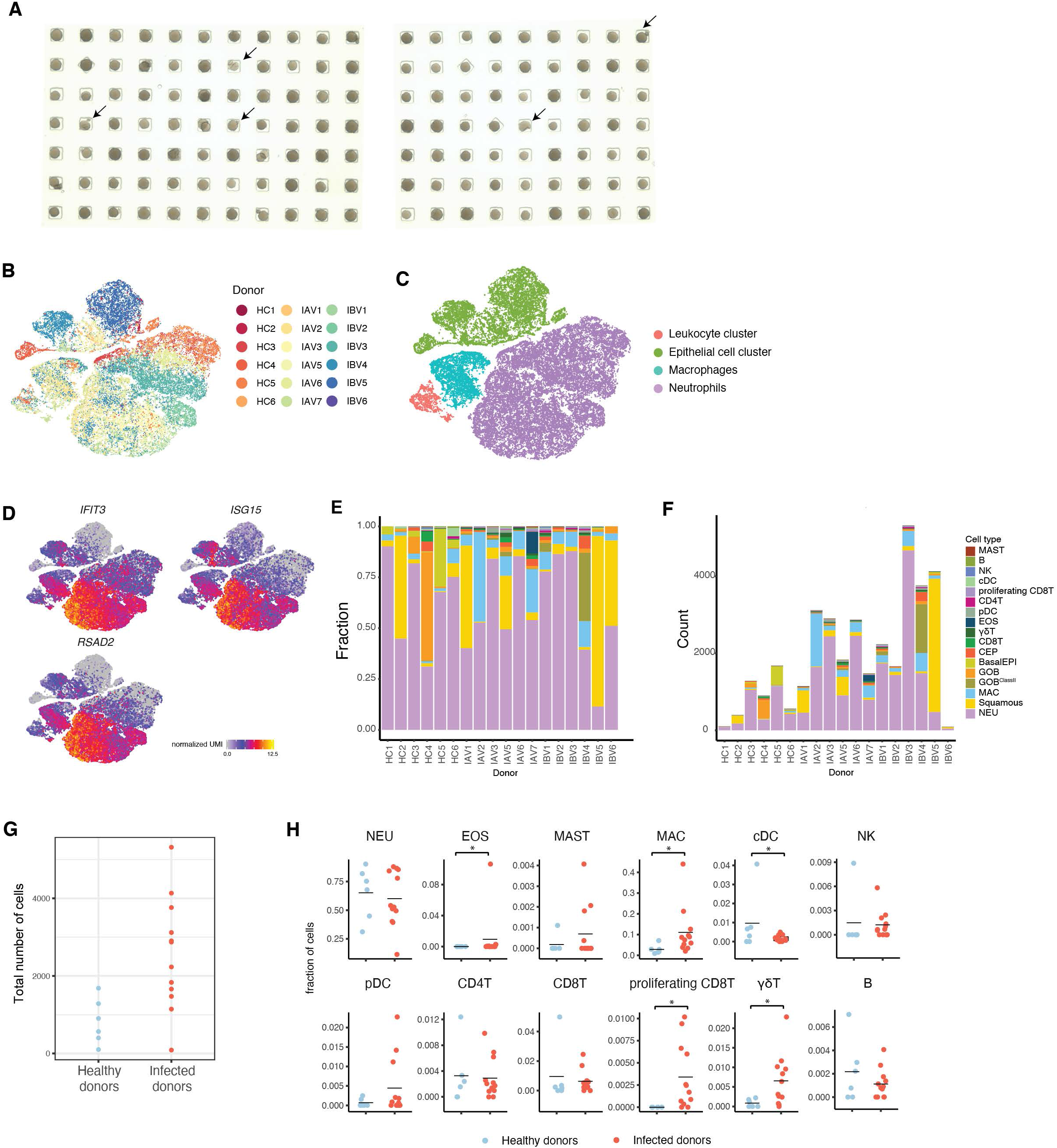
Related to Figure 1 Cell type distributions across donors. **A.** Images of bead loading on Seq-Well platform. Arrows point to the speculated broken beads. **B**. tSNE of all cells colored by donors. **C.** tSNE of all cells colored by the four major cell type clusters. **D.** Normalized expression of three interferon stimulated genes (*IFIT3, ISG15, RSAD2*). **E**. Fraction of cell types across donors. **F.** Raw counts of each cell type for each donor. **G.** The total number of cells in scRNA-Seq data collected from healthy, IAV- or IBV-infected donors. The total number of cells collected from influenza-infected donors is significantly increased compared to those from healthy donors (t-test, *P* =0.003). **H.** Fraction of each immune cell type in infected or healthy donors. The number of each cell type from each healthy (n=6) and infected (n=12) donor is plotted here. The black line shows the mean. *: p < 0.05, Wald test.

**Figure S2.**
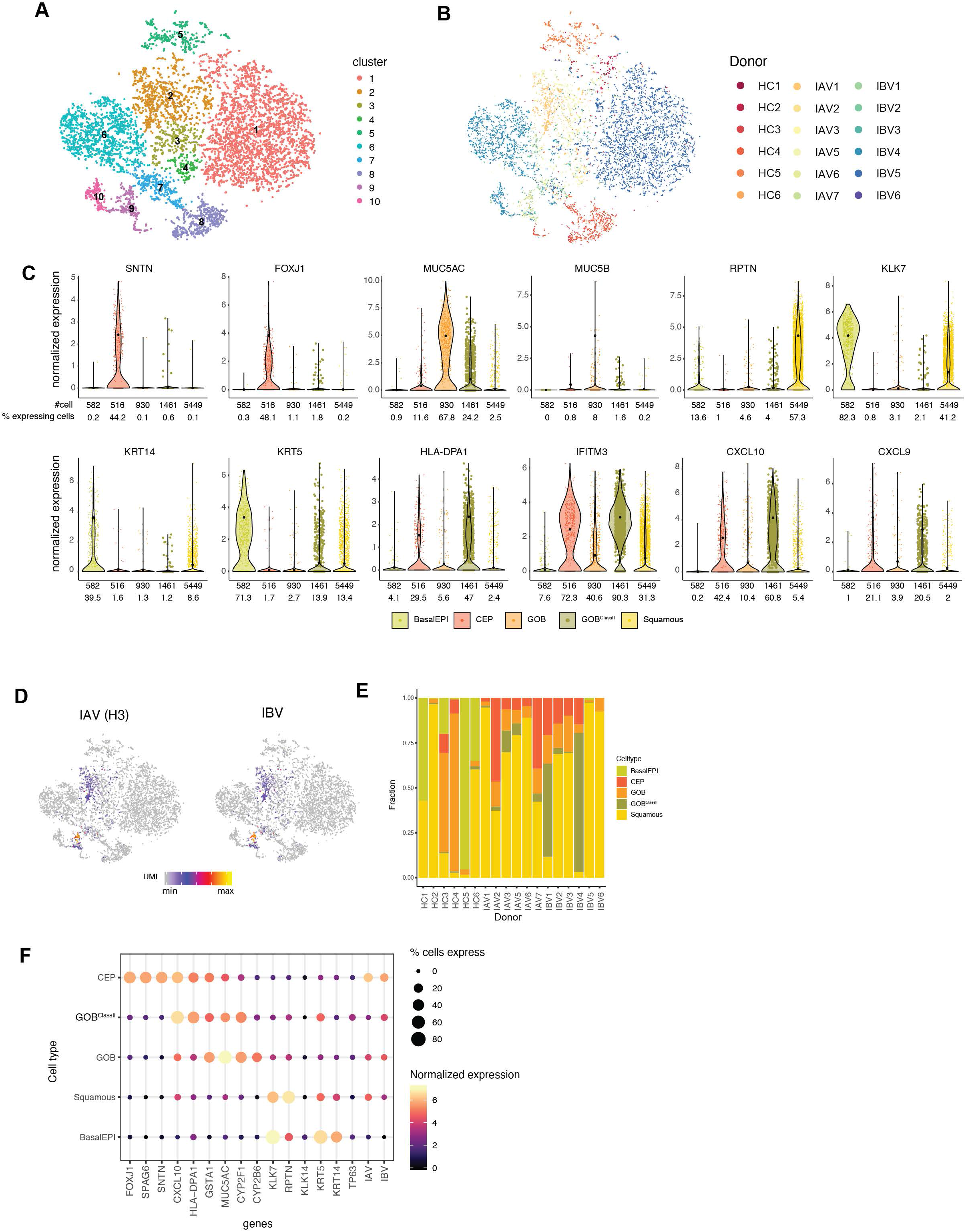
Related to Figure 1 Epithelial cell cluster cell type identification. **A.** Ten clusters were called by density clustering from the epithelial cell cluster. **B.** tSNE plot colored by donors. **C.** Normalized expression of cell type markers in each cell type. The black points are the mean expression of expressing cells. **D.** tSNE plots on the influenza viral transcript expressions. **E.** Fraction of epithelial cell types found in each donor. **F**. The expression levels (normalized expression and percentage of cells expressing) of marker genes per epithelial cell type.

**Figure S3.**
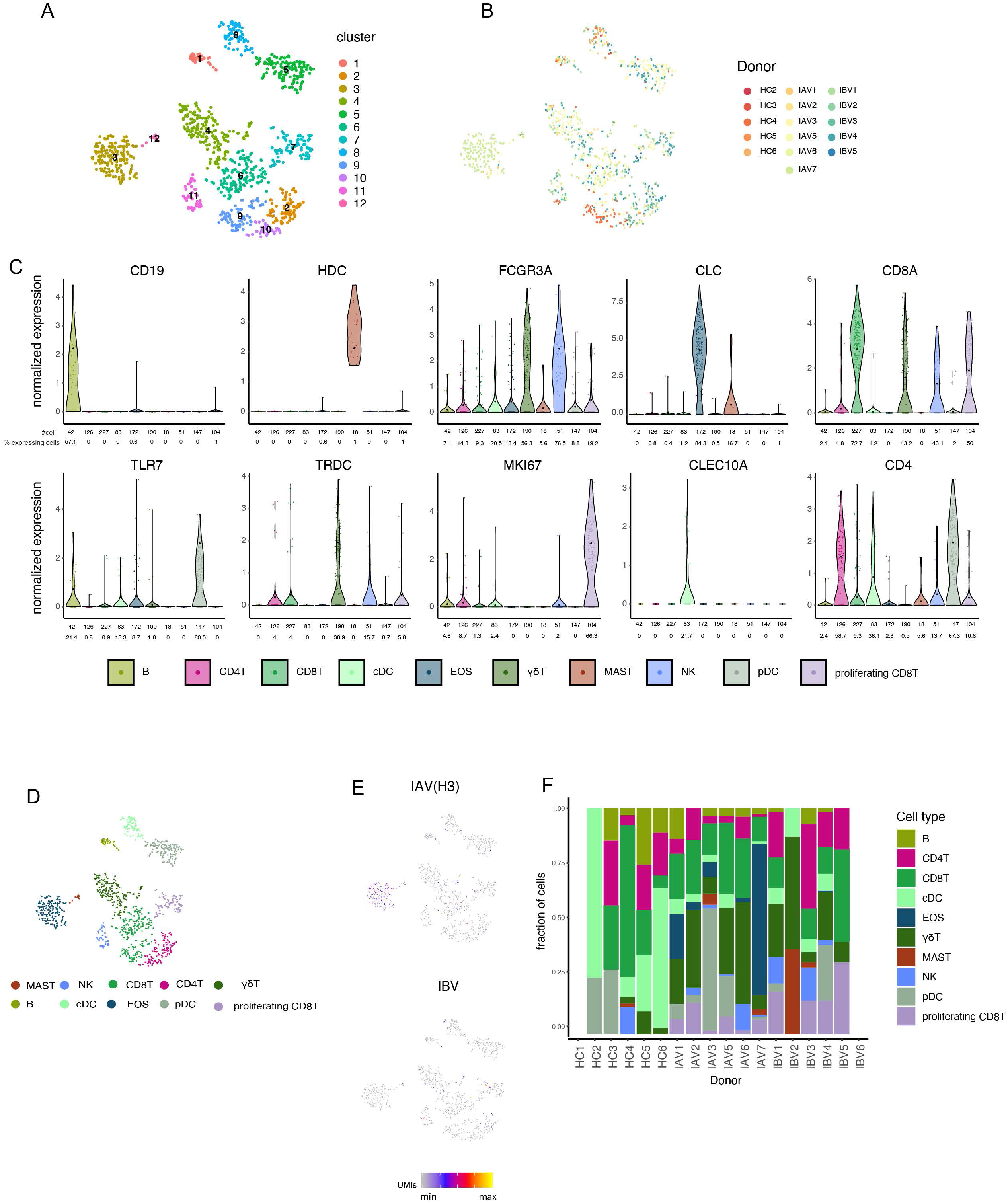
Related to Figure 1 Leukocyte cluster cell type identification. **A.** Twelve clusters were called by density clustering from the leukocyte cell cluster. **B.** tSNE plot colored by donors. **C.** Normalized expression of cell type markers for each cell type in the leukocyte cluster. The black points are the mean expression of expressing cells. **D.** Cell types identified. **E.** tSNE plots on the influenza viral transcript expression. **F.** Cell types found in each donor. HC1 and IBV6 do not yield cell types shown here.

**Figure S4.**
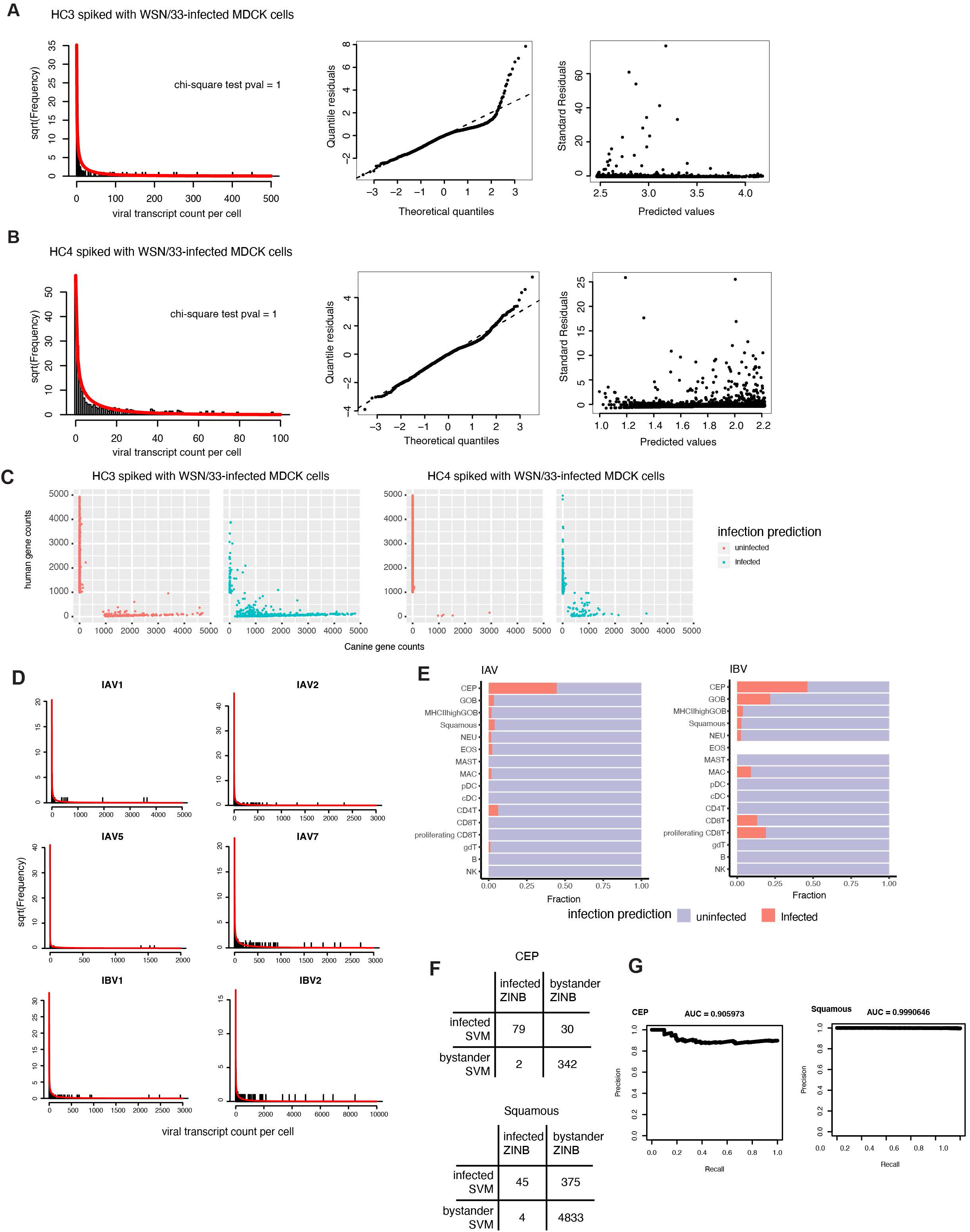
Related to Figure 2 Background viral transcript estimation and infected cell type identification. **A.** Histogram of viral counts in human nasal wash cells from HC3 nasal wash cells spiked-in with WSN-infected MDCK cells. The red line is the fit of viral counts predicted with the ZINB model. **B.** histogram of viral counts in human nasal wash cells from HC4 nasal wash cells spiked-in with WSN-infected MDCK cells. The red line is the fit of viral counts predicted with the ZINB model. **C.** The expression of human cells and dog genes in each cell in HC3 and HC4 nasal wash cells spiked-in with WSN-infected MDCK cells. Cells are colored by their predicted infection state. **D.** Histogram of viral counts in each of the highly infected donors. The red line is the fit of viral counts predicted with the ZINB model. **E.** Fraction of cells predicted to be infected by ZINB models for each cell type, broken down by cells from either IAV- or IBV-infected donors. **F.** Infected and bystander cells predicted by ZINB models and classified by SVM for CEP and Squamous cells. The result of SVM classification for CEP and Squamous cells was used for infected cell classification. **G.** Precision-Recall curve for each SVM classifier.

**Figure S5.**
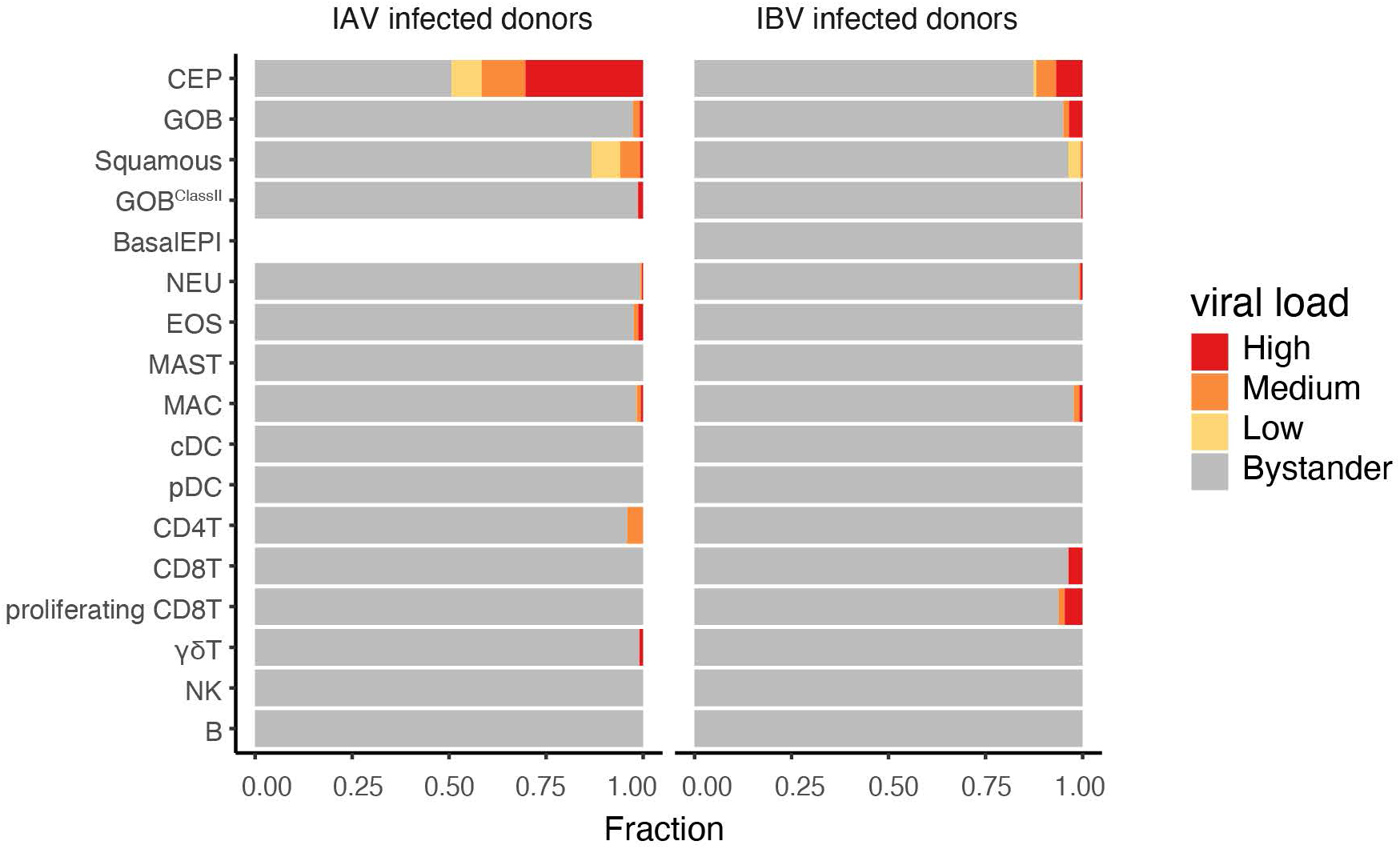
Related to Figure 2 Viral load states breakdown in IAV- or IBV-infected donors. Fraction of cells classified in each viral load state for cells from IAV- or IBV-infected donors

**Figure S6.**
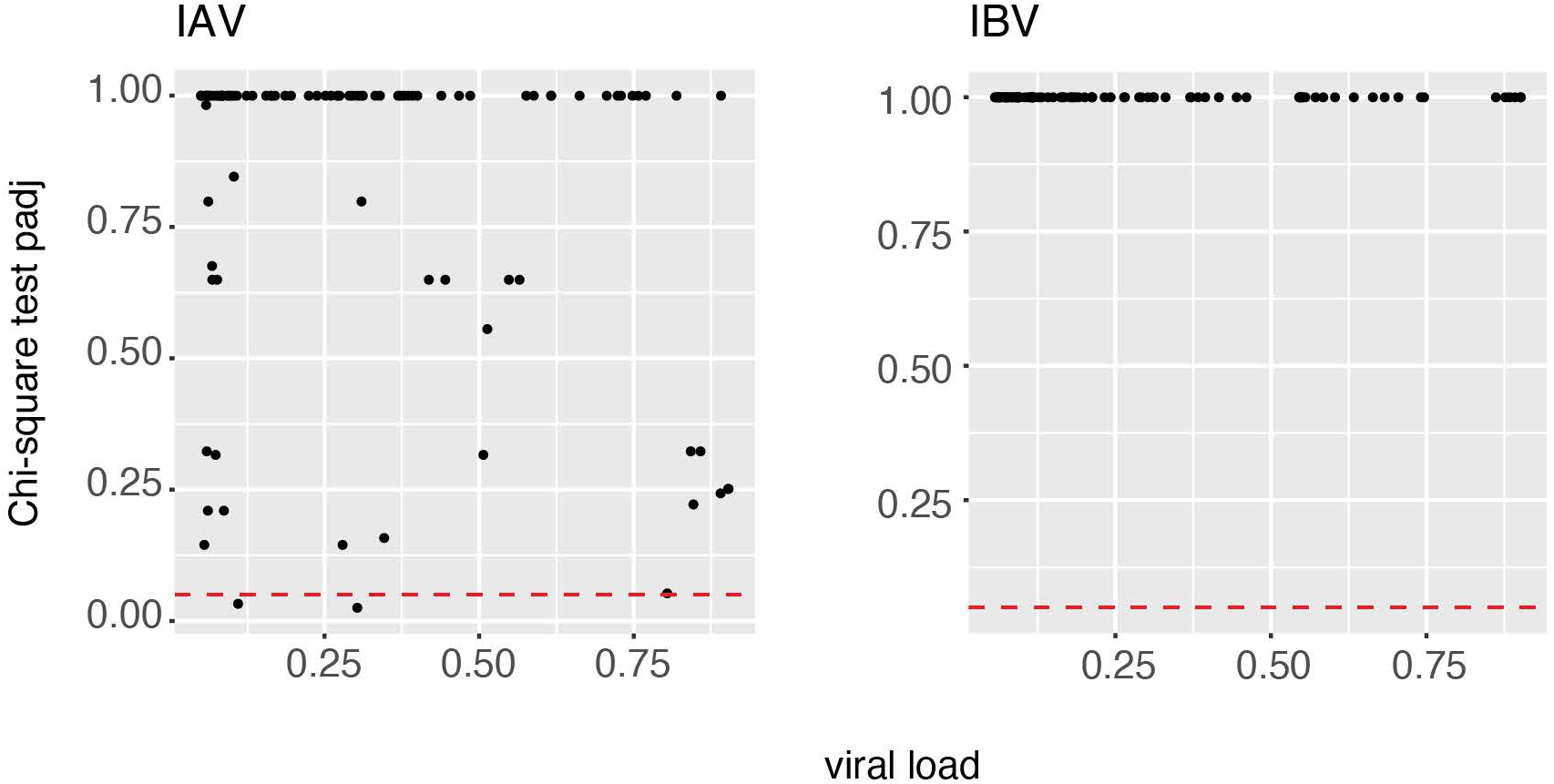
Related to Figure 3 IAV segment expression is more variable. Chi-squared test was applied to each IAV and IBV high viral load cell to test if the expression of each viral genome segment is equally likely. The adjusted p-values are plotted against the viral load of each cell. The red dash line intercepts the y axis at 0.05.

**Figure S7.**
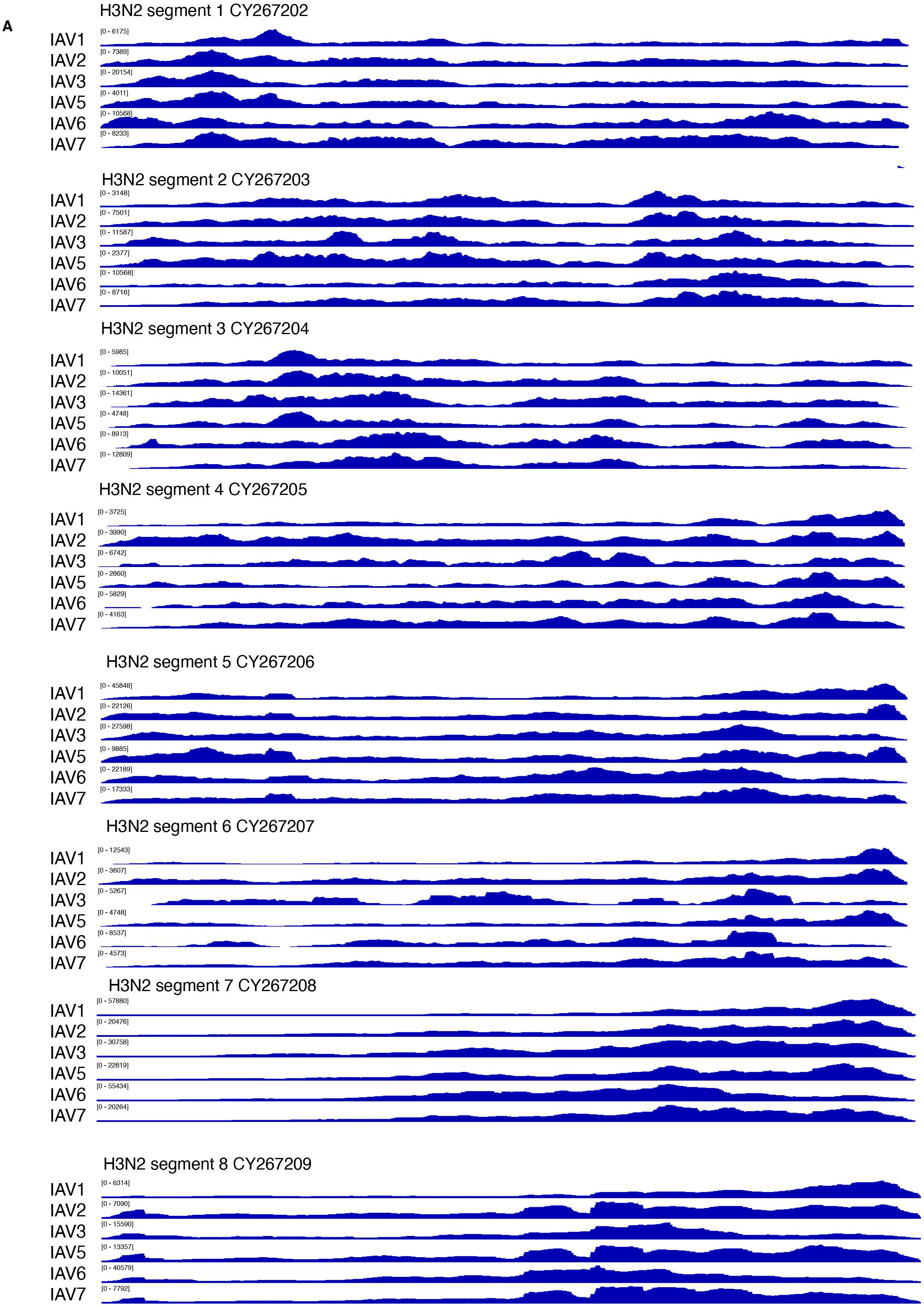
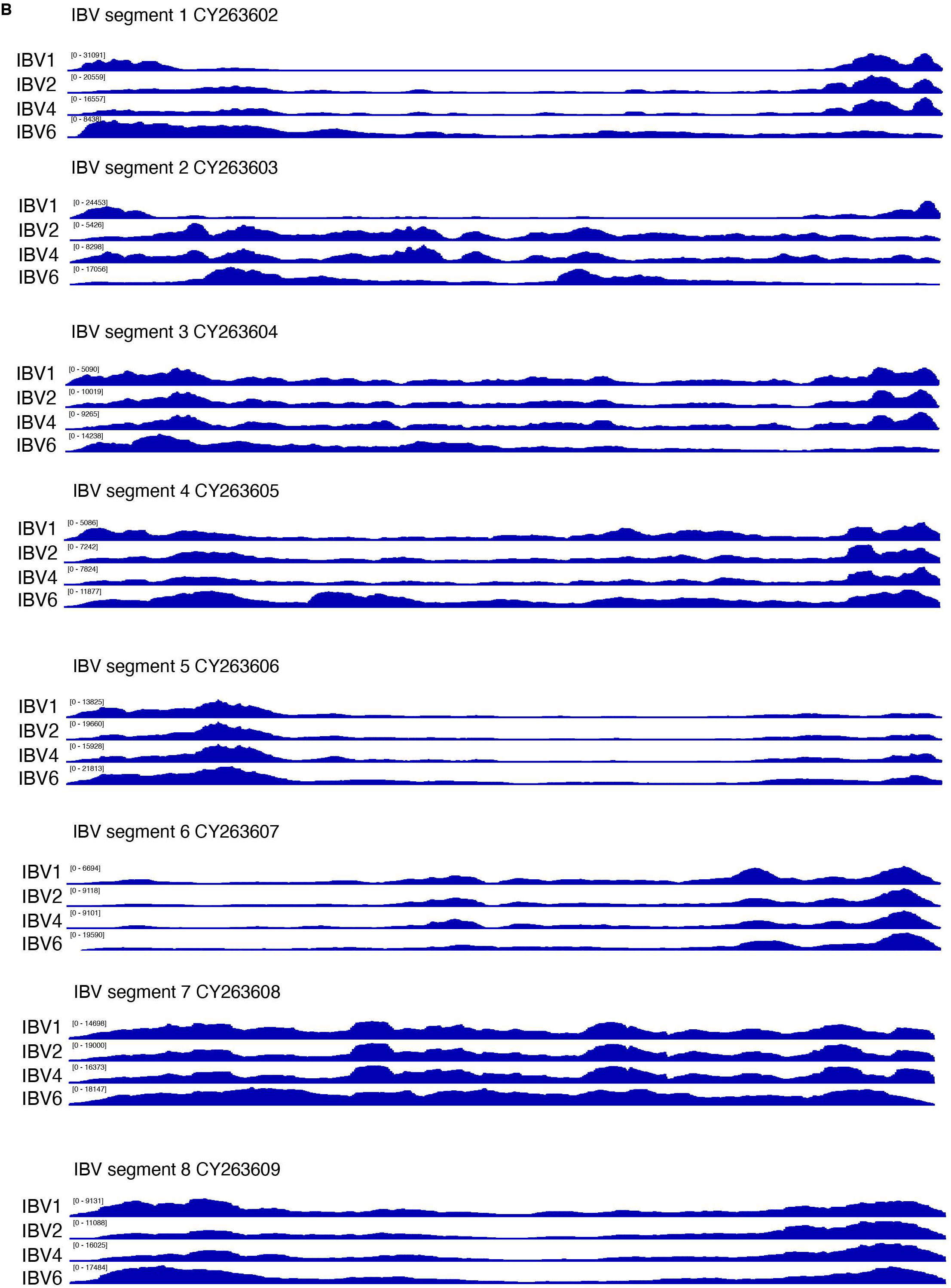
Related to Figure 4 Read coverage on influenza genomes from the scRNA-Seq data for all twelve influenza virus-infected donors. Reads from scRNA-Seq libraries mapped to each viral genome are plotted here. The minimum and maximum numbers of reads covering each base are noted. PCR duplicated reads were removed. The average PCR duplication rate is estimated to be 4x. **A.** Read coverage for IAV infected donors. **B.** Read coverage for IBV infected donors.

**Figure S8.**
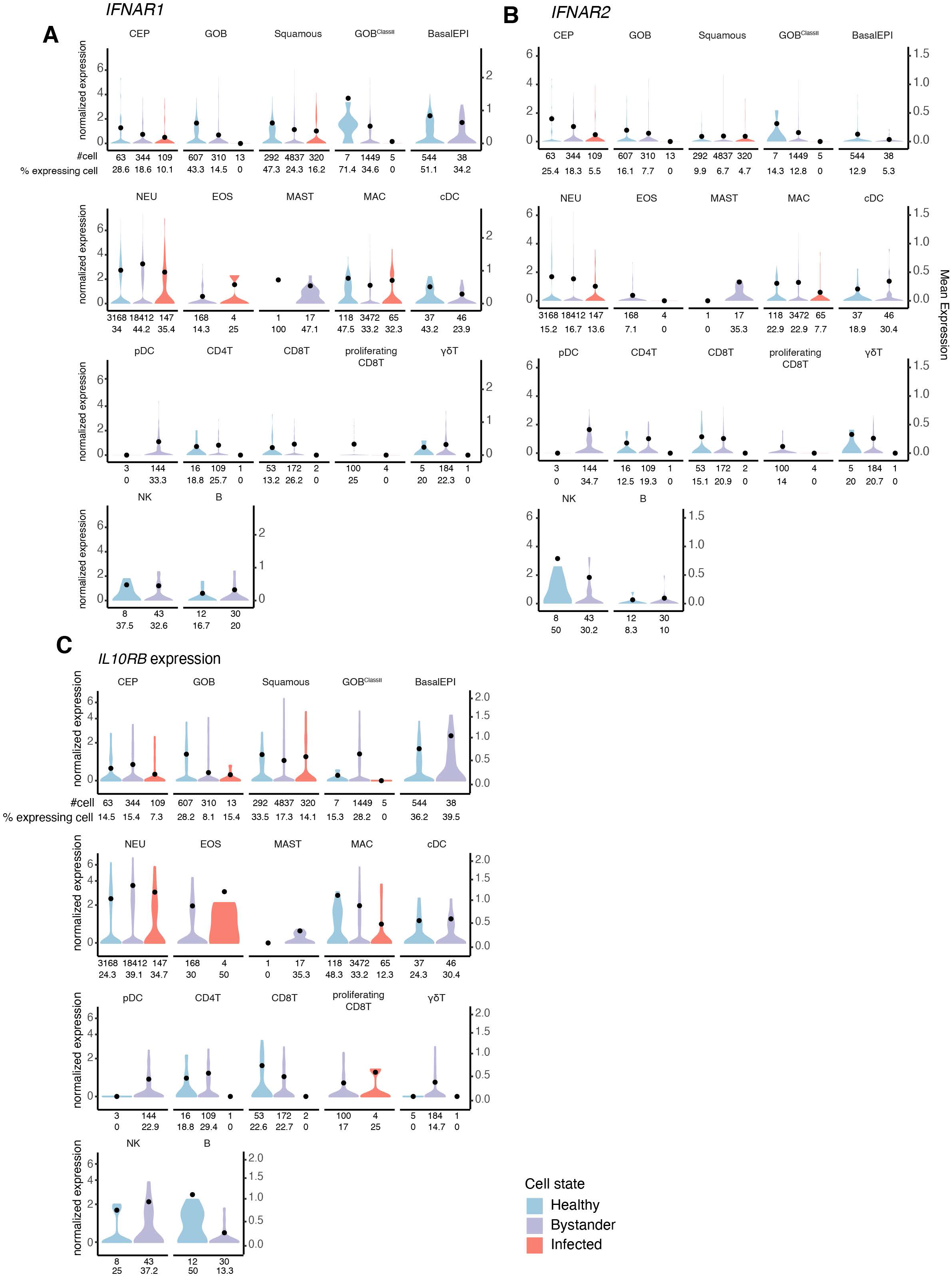
Related to Figure 5 Expressions of type I and III IFN receptors in different cellular states across cell types. Violin plots of the normalized expressions of *IFNAR1*, *IFNAR2*, and *IL10RB* in different cellular states across cell types. The black points are the mean expression of expressing cells. **A.***IFNAR1* normalized expressions. **B.***IFNAR2* normalized expressions. **C.***IL10RB* normalized expression.

**Figure S9.**
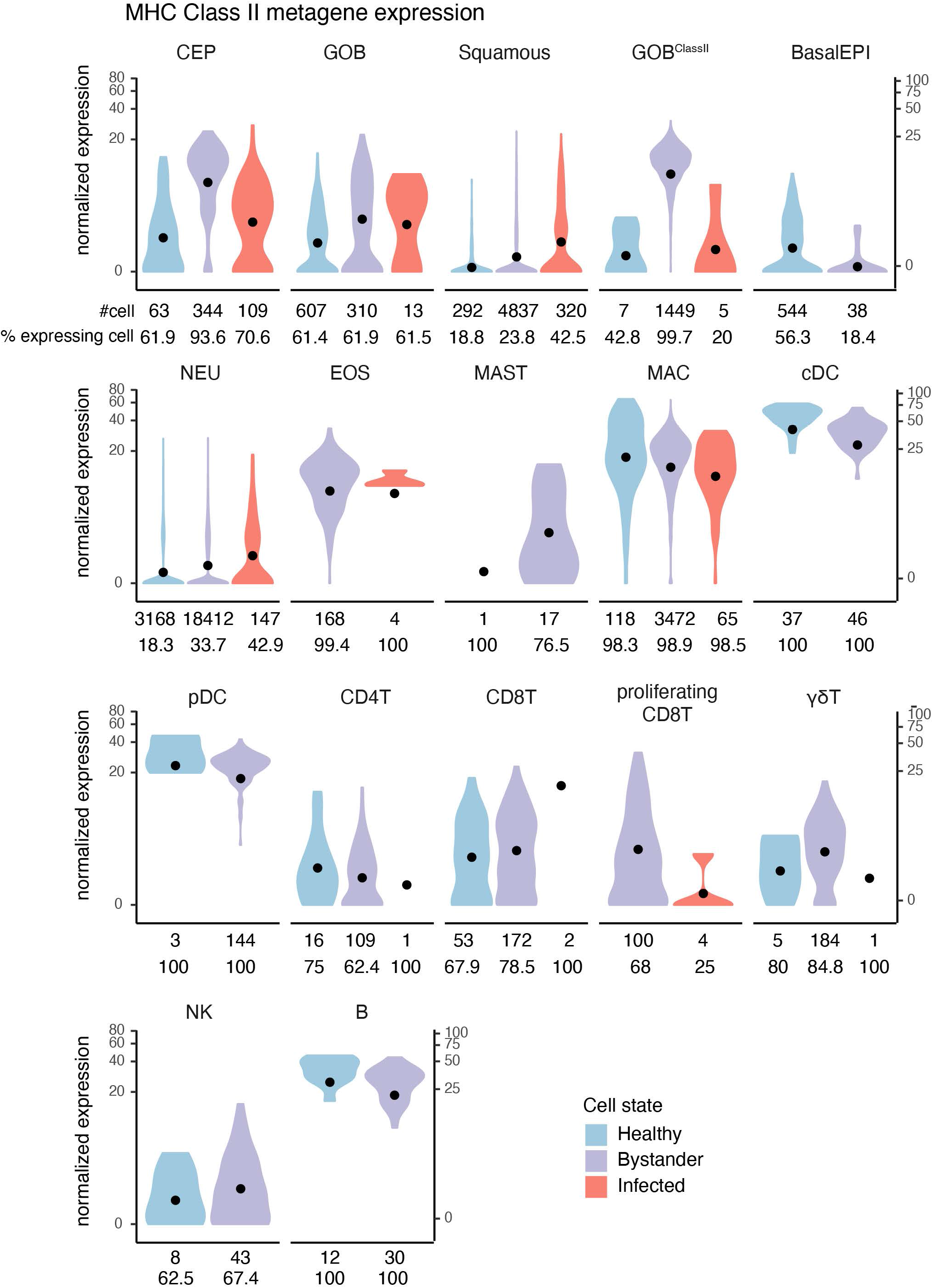
Related to Figure 5 MHC class II gene expression in different cellular states across cell types. The normalized expression of MHC class II metagene is calculated by the sum of the normalized expression of all MHC class II genes mapped in the scRNA-Seq data and *CIITA,* the gene encoding the transcription factor for MHC class II. The black points are the mean expression of MHC class II metagene.

## Methods

### Sample collection

#### Approvals from Institutional Review Boards

All procedures were approved by the University of Massachusetts Medical School Institutional Review Board (IRB protocol # H00009277) and participants signed an informed consent document whenever required by IRB.

#### Subject enrollment

Nasal washes were obtained from adult healthy controls and from adults with diagnosis of acute influenza A or B by rapid antigen test (Flu A or B antigen, direct fluorescence antigen test) and/or by respiratory virus panel (PCR testing for influenza A, influenza A H1, influenza A H3, influenza B, adenovirus, metapneumovirus, respiratory syncytial virus A, respiratory syncytial virus B, rhino/enterovirus, parainfluenza 1, parainfluenza 2, parainfluenza 3), who show symptoms up to seven days. Samples were obtained by irrigation of each naris with up to 10 mL of saline, and collected in a single container. The sample was then transported to the research laboratory for processing. Upon receipt, the sample was immediately stored on ice and 10 mL cell growth media (DMEM or RPMI1640 with 10% fetal bovine serum) was added. The material was strained using a 40 μM nylon cell strainer (Corning) into a 50 mL centrifuge tube. Cells were pelleted at 1300 rpm for 10 min at 4°C. All but 1 mL of supernatant was discarded, the pellet resuspended in the remaining 1 mL of supernatant, and material was transferred to an Eppendorf tube and pelleted at 2000 rpm for 5 min. If the pellet contained visible blood, 200 μL of RBC lysis solution (Sigma) was added to resuspend the pellet and incubated at room temperature for 2 min, after which 1 mL of cell media was added, and the cells were pelleted at 2000 rpm for 5 min. The final pellet was resuspended in up to 1 mL of media and quantified.

For two healthy donor samples (“HC3” and “HC4”), 2,000 influenza A/WSN/33-infected MDCK cells (infected for overnight at a multiplicity of infection of 1) were added to 18,000 primary human cells prior to loading on the Seq-Well array. This was performed to measure any potential cross-contamination from dead cells and free viral transcripts in the sample. Association of either canine or A/WSN/33 viral transcripts with human cell barcodes, or vice versa, would suggest that a well contained both human and canine material (i.e., either multiple cells per well or RNA contaminants from lysed cells are present).

Influenza A/WSN/33 was passaged in MDCK cells (MDCK ATCC catalog number: PTA-6500. A/WSN/33 ATCC catalog number: VR-825).

### RNA sequencing

#### Seq-Well

Seq-Well was used according to published methods (Gierahn et al., 2017) to capture single cells on a microwell array. Each microwell has only one bead carrying oligonucleotides that have a cell barcode, unique molecular identifiers (UMIs), and a polyT tail. Each array was loaded with 20,000 cells. Any remaining cells were put in TRIZOL and stored at −80°C. After cell lysis, mRNA transcripts were captured by the oligonucleotides on the bead. The cDNA libraries were prepared using Illumina Nextera XT Library Prep Kits and sequenced using the NovaSeq 6000x System from Illumina. NovaSeq S2 100-cycle sequencing was performed at the Broad Institute Genomics Platform.

#### Bulk RNA Sequencing

Bulk RNA-Seq was performed for a subset of the samples initially stored in TRIZOL® Reagent, Cat.:15596. RNA extraction was performed following the manufacturer’s directions, and libraries were constructed using the NuGen Ovation FFPE RNA-Seq Multiplex System. Libraries were sequenced on an Illumina NextSeq 500.

#### RT-qPCR

One step RT-qPCR was performed with TaqMan chemistry. Primers and probes used for IAV-M, IBV-HA were previously reported (World Health Organization, 2017) and synthesized by Bio-Rad Laboratories. B2M primer and probe were made by ThermoFisher Scientific (Cat #: 4326319E) and RNA samples were run in triplex with IAV and IBV. RT-qPCR amplification was carried out in 10 μL reactions and 3 replicates were run for each sample. Each plate contained a range of serial dilutions of viral RNA from B/Mass/3/66 (ATCC, Cat #: VR-523), viral RNA from A/PR/8/34 (Charles River Laboratories) and A549 cellular RNA for standard curve generation. The QuantiFast Pathogen RT-PCR+IC Kit from Qiagen (Cat #: 211452) was used. Experiments were conducted to test inter- and intraplate variability. The real-time PCR amplification was performed on a CFX96 C1000 thermal cycler (Bio-Rad Laboratories) with the following condition: 50°C for 20 min, 95°C for 5 min, 40 cycles of 95°C for 15 sec and 60°C for 45 sec. The results were analyzed with built-in software. The ratios of IAV-M/B2M, IBV-HA/B2M were calculated using standard curves.

### Computational Analysis

#### Genome Alignment

Reads were aligned to the GRCh37 reference genome combined with influenza genomes. A/Massachusetts/20/2017 (H3N2): genome ID: CY264272, CY264273, CY264274, CY264275, CY264276, CY264277, CY264278, CY264279; B/Massachusetts/18/2017 (Yamagata lineage): genome ID: CY263602, CY263603, CY263604, CY263605, CY263606, CY263607, CY263608, CY263609. Reads were additionally aligned to a panel of respiratory virus genomes: Respiratory syncytial virus A (RSVA), RSVB, human parainfluenza virus, human respirovirus, rubulavirus, mumps virus, human rhinovirus (HRV) A, HRVB, HRVC, human adenovirus B2 and influenza C virus with the genome IDs: KJ643560, KC283039, KY674966, KY674953, KY967354, KY779616, KY674950, MF965239, KY369875, KY369902, KY369880, NC011202, KM504277, KM504278, KM504279, KM504280, KM504281, KM504282, KM504283. Mapped reads from each sample were then corrected for DropSeq barcode synthesis error using the DropSeq core computational tools developed by the McCarroll Lab (Macosko and Goldman). Genes were quantified using End Sequence Analysis Toolkit (ESAT, github/garber-lab/ESAT) with parameters *-wlen 100 -wOlap 50 -wExt 0 -scPrep* (Derr et al., 2016). Finally, UMIs that likely result from sequencing errors were corrected by merging any UMIs that were observed only once and have 1 hamming distance from a UMI detected by two or more aligned reads.

Only cell barcodes with more than 1,000 UMIs were analyzed. Cell barcodes with mostly erythrocyte genes (*HBA, HBB*) were removed. From here on, the remaining cell barcodes in the matrix would be referred to as cells. The final gene by cell matrix was normalized using the scran package v3.10 (Lun et al., 2016). The normalized matrix was used for dimensionality reduction by first selecting variable genes that had a high coefficient of variance (CV) and were expressed (>=1 UMI) by more than three cells. Influenza viral genes, interferon stimulated genes, and cell cycle related genes were removed from the variable gene list in order to minimize the impact of viral responses and mitosis on clustering and cell type identification. This resulted in the selection of 2484 variable genes. t-distributed stochastic neighbor embedding (tSNE) was applied to the first ten principal components (PCs), which explained 95% of the total data variance (Maaten and Hinton, 2008).

#### Clustering and cell type identification

Density clustering (Rodriguez and Laio, 2014) was performed on the resulting tSNE coordinates (**Figure 1B,C**) and identified four major clusters: epithelial cells, neutrophils, macrophages and leukocytes (**Figure S1E**). The epithelial cell cluster and the leukocyte cluster were then re-clustered independently, as described above, to identify populations within each metacluster. Specifically, the epithelial cell cluster was re-embedded using 2629 variable genes selected by the same criteria mentioned in the previous section and 13 PCs that explained 95% of the variance. Density clustering on the epithelial cell tSNE map (**Figure S2A**) revealed ten clusters. Differential gene expression analysis using edgeR (Robinson et al., 2010) was performed to identify marker genes for each cluster (**Figure S2C**). For the leukocyte metacluster, 2583 variable genes were selected by the same criteria, and 8 PCs that explained 90% of the variance were used as input to tSNE. In order to identify specific T cell populations, the following procedures were followed: cells were clustered based on the tSNE map into six clusters. The top 300 differentially expressed genes (DEGs) that distinguish each of the six clusters from all other leukocytes were selected, resulting in 1454 unique genes after removing viral transcripts and interferon responding genes. This matrix of 1454 genes by 1160 cells was used for dimensionality reduction through PCA and tSNE, followed by density clustering. The second round of clustering revealed different T cell groups, including CD4+ T cells, CD8+ T cells, proliferating CD8+ T cells, and γδT cells, and a small population of B cells and mast cells.

#### Identifying viral infected cells

To estimate a sample-specific distribution of ambient influenza mRNA contamination from cell lysis and identify truly infected cells, two control libraries were generated by spiking-in influenza A/WSN/33-infected Madin-Darby canine kidney (MDCK) cells to nasal wash cells from two healthy human donors prior to performing scRNA-Seq. These libraries enabled tracking of reads mapped to each organism and estimation of the amount of ambient RNAs per cell.

Sequencing reads from WSN-infected MDCK spiked-in donors were mapped to a hybrid genome of GRCh37 reference genome, camFam 3.1, and A/WSN/33 influenza genome (CY034132, CY034133, CY034134, CY034135, CY034136, CY034137, CY034138, CY034139). The mapped reads were then processed through the same pipeline described in **Genome Alignment.** Only cells with > 1000 UMIs were analyzed. Human cells and MDCK cells formed distinct clusters. MDCK transcripts and viral transcripts were found in human cells, which suggested those transcripts were the result of ambient RNA contamination. The distribution of ambient viral RNAs was overdispersed with zero inflation. We also observed that the viral transcripts and the total number of UMIs per cell are correlated. To assess the amount of ambient RNA in the cells, a zero-inflated negative binomial model (ZINB) was built using the function hurdle (package “pscl v1.5.2”) in R (v3.5.0). The observed viral counts were modeled against the total number of UMIs in each cell for each spiked-in donor. The model performance was evaluated using Q-Q plots and residuals. This model was only applied to highly infected donors (IAV1, IAV2, IAV5, IAV7, IBV1, IBV2) to assess the expected fraction of viral ambient RNA per cell.

It is notable that human cells carrying high levels of A/WSN/33 transcripts were predicted to be infected after applying the ZINB model (**Figure S4C-D**). Since only a small fraction of these cells also have a high fraction of dog mRNAs, human/dog doublets do not explain the high presence of influenza mRNAs. This suggests that they may have been infected during the time in which they were mixed together (∼1.5hr) before cell lysis on the Seq-Well platform. Indeed, a disproportionate number of cells with high influenza viral mRNAs are epithelial (170 epithelial cells and 64 immune cells), the natural target of influenza virus, further indicating that infection after mixing occurred.

The ZINB model had no power to predict infection in the case of cells with lower levels of viral transcripts. This includes the 298 virus positive cells from half of the infected donors, and a large number of virus positive cells that were not predicted to be infected in high viral load donors. To ensure that infected cell identification was robust, a support vector machine (SVM) classifier was applied to distinguish between infected and bystander cells. The SVM classifier was only applied to cell types which had more than 10 cells predicted to be infected by the ZINB model, namely CEP and Squamous cells. The gene expression signature related to viral infection for each cell type was generated by comparing infected cells predicted by ZINB and bystander cells with no viral transcripts using *edgeR* (Robinson et al., 2010). Genes expressed by more than 20% of the cells were tested, and genes with at least a 2 fold change (FDR < 0.05) in infected cells predicted by ZINB were selected as features for the model. The SVM classifier for each cell type was trained using 70% of the infected cells predicted by ZINB and an equal number of bystander cells without any viral transcripts. The remaining cells from each cell type were then classified as infected or bystander cells. The SVM classification for CEP and Squamous cells showed 3% and 6% false positive rates, respectively.

#### Cell type enrichment statistical test

The method described previously by (Xu et al., 2019) was followed to test for cell type composition changes between healthy and infected donors. Briefly, a negative binomial regression model was used to assess if the cell type composition between healthy donors and infected cells are significantly different. For each cell type, the infection label (healthy or infected) was used as a covariate and the total number of cells sampled from each donor was used as an offset variable. The Wald test was used on the regression coefficient to assess the P value for each cell type.

#### Differential gene expression analysis

Differential gene expression analysis was performed using *edgeR,* comparing bystander cells to healthy cells, and infected cells to bystander cells for each cell type. Comparisons with fewer than 3 cells in any cellular infection state were not tested. To generate cell type markers, *edgeR* was used to test for differentially expressed genes of each cell type against the rest sixteen cell types with equal weights. Genes were considered differentially expressed with at least 2 fold change and false discovery rate less than 0.05.

#### Gene Set Enrichment Analysis (GSEA)

To capture more subtle differences among cell states which affect functionally related genes, gene set enrichment analysis (GSEA v3.0) (Subramanian et al., 2005) was performed using a ranked gene list constructed from the *edgeR* results. Genes were ranked by their reported fold change (logFC). GSEA was run via command line using parameters *--nperm 2000 -set_max 3000*. The GO term annotation reference was obtained from MSigDB C5 collection (http://software.broadinstitute.org/gsea/msigdb/genesets.jsp?collection=C5). Significantly enriched gene ontology (GO) terms were selected based on (1) the normalized enrichment score (NES > 2 or NES < −2), and (2) the false discovery rate (FDR < 0.05).

#### Aggregated pseudobulk gene expression calculation

The aggregated pseudobulk gene expression was calculated as ∑g_(1,2…i)_/∑c_(1,2…i)_ * 1,000,000 for all i cells in a group, where g is the normalized UMI count of a gene, and c is the total number of UMI counts in a cell.

#### Virus genotyping and single nucleotide variants (SNVs) calling

For each donor’s scRNA-Seq library, reads mapped to influenza viral genomes were removed of PCR duplicates using *umitools -dedup (Smith et al.)*. The remaining reads were then used to identify SNVs using mpileup from samtools via command line (Li, 2011). SNVs were called in each donor with following criteria: for a highly covered position (> 20 reads), a SNV is called when more than 50% of mapped reads support the alternative allele; for a lowly covered position (≤ 20 reads), if the alternative allele is supported by > 50% of the reads and this allele was also found in other donors with higher coverage, a SNV will be called, otherwise this position will not be called in this donor. SNVs called from the scRNA-Seq data were also detected using the bulk RNA-Seq data for donors who have both types of library, confirming the accuracy of the SNV detection approach in scRNA-Seq data. A hamming distance matrix of the viral sequence in each donor was calculated by using only the SNV positions covered in all donors. Hierarchical clustering was used to identify the similarity of the viruses among donors. **Supplemental Table 2** contains details on the SNV called per donor and per viral genome segment.

### Special notes: Bead breakage correction

After single-cell mRNA capturing on the Seq-Well array, cells from each donor were split into five aliquots, in order to control the number of cells to sequence from each sample. In theory, the cell barcodes within aliquots from each donor should collide minimally (1.3% for 78,000 beads determined from random 11 bp oligonucleotide simulation). However, we observed the same barcode in different aliquots from the same sample, at a much higher rate than expected from random barcode sampling. Further analysis on these barcodes revealed that they had similar transcriptome profiles. Based on light microscopy inspection, we hypothesized that this phenomenon could result from the broken beads that were separated into different aliquots. This “bead breakage problem” was corrected by merging cell barcodes across aliquots from the same donor with non-random barcode collision whenever they exhibit similar transcriptome profiles.

### Special notes: Illumina sequencing index swapping correction

Index swapping (hopping), i.e., the integration of free adapters, happens on Illumina patterned flow cells for sequencing platforms such as NovaSeq, HiSeq4000 and HiSeqX(Costello et al., 2018). This issue impacted our analysis when identifying infected cells, genotyping the virus sequences and calling SNVs. We estimated the index swapping rate for each flow cell by identifying the same cell barcodes. The swapped reads had the same barcode as cell barcodes from another sample. Since Seq-Well utilizes Drop-Seq beads, which have the 12 bp cell barcodes randomly generated by split-and-pool, random barcode collision can happen. We estimated the rate of random barcode collision to be 1.3%, given 78,000 beads loaded on each Seq-Well array. We also aliquoted each sample into five and each aliquot was indexed differently, so the aforementioned bead breakage problem made estimation of the swapping rate and identification of swapped reads challenging. To identify the swapped reads from each sample, we only focused on the cell barcodes with more than 1000 UMI (i.e., true cells) and identified the same barcodes from other samples on the same flow cell. We removed the reads with those barcodes from other samples in cases where the amount corresponded to less than 5% of all reads with the same barcodes on the same flow cell.

## Acknowledgments

This work was supported by the UMass Center for Clinical and Translational Science Project Pilot Program, the Office of the Assistant Secretary of Defense for Health Affairs, through the Peer Reviewed Medical Research Program (award no. W81XWH-15-1-0317) and the Richard and Susan Smith Family Foundation (J.O-M).

We thank multiple individuals at UMass Medical School, including Dr. Gregory Pazour for guidance in identifying and classifying ciliated cells and cell markers, Drs. David Paquette and Melanie Trombly for assistance with manuscript submission and for providing feedback throughout the project, Jaclyn Longtine, Karen Longtine and Melissa O’Neill for sample collection, Alan Derr, Jake Gellatly, Pyae Kwaye, Maureen Hester and Elham Ahmadi for data analysis discussions. We also thank Travis Hughes and Marc Wadsworth (Shalek lab, M.I.T.) for advice on Seq-Well.

## Author contributions

R. W. F., J. P. W., and M. G. conceived and designed the study, advised on data collection and supervised the research. Y. C. designed and performed bioinformatics analysis under the guidance of M. G.; Z. G. prepared and processed all samples for scRNA-Seq and bulk RNA-Seq, with supervision by J. O.-M., P. McD., M. G., R. W. F., and J. P. W.; P. V. and Y. C. designed and applied the ZINB model for identifying infected cells; Y.C. and E.D. built and applied the SVM classifier for infected cell classification on epithelial cells; P. L. performed qPCR experiments under J. P. W.’s guidance. All authors participated in data interpretation. Y. C., R. W. F., J. P. W., and M. G. wrote the manuscript with input from all authors.

